# Atoh1 is required for the formation of lateral line electroreceptors and hair cells, whereas Foxg1 represses an electrosensory fate

**DOI:** 10.1101/2023.04.15.537030

**Authors:** Martin Minařík, Alexander S. Campbell, Roman Franěk, Michaela Vazačová, Miloš Havelka, David Gela, Martin Pšenička, Clare V. H. Baker

## Abstract

In electroreceptive jawed fishes and amphibians, individual lateral line placodes form lines of neuromasts on the head containing mechanosensory hair cells, flanked by fields of ampullary organs containing electroreceptors - modified hair cells that respond to weak electric fields. Extensively shared gene expression between neuromasts and ampullary organs suggests that conserved molecular mechanisms are involved in their development, but a few transcription factor genes are restricted either to the developing electrosensory or mechanosensory lateral line. Here, we used CRISPR/Cas9-mediated mutagenesis in F0-injected sterlet embryos (*Acipenser ruthenus*, a sturgeon) to test the function of three such genes. We found that the ‘hair cell’ transcription factor gene *Atoh1* is required for both hair cell and electroreceptor differentiation in sterlet, and for *Pou4f3* and *Gfi1* expression in both neuromasts and ampullary organs. These data support the conservation of developmental mechanisms between hair cells and electroreceptors. Targeting ampullary organ-restricted *Neurod4* did not yield any phenotype, potentially owing to redundancy with other *Neurod* genes that we found to be expressed in sterlet ampullary organs. After targeting mechanosensory-restricted *Foxg1*, ampullary organs formed within neuromast lines, suggesting that Foxg1 normally represses their development. We speculate that electrosensory organs may be the ‘default’ fate of lateral line primordia in electroreceptive vertebrates.

## Introduction

Mechanosensory hair cells in different compartments of the inner ear transduce fluid movements for hearing and balance (Moser et al., 2020; Caprara and Peng, 2022; Mukhopadhyay and Pangrsic, 2022). In fishes and aquatic-stage amphibians, hair cells are also found in lateral line neuromasts in the skin, which are stimulated by local water movement (Mogdans, 2019; Webb, 2021). Supporting cells in neuromasts differentiate into hair cells for homeostasis and repair (Kniss et al., 2016; Lush et al., 2019) and neuromasts are easily accessible in lines over the head and trunk, making the zebrafish lateral line an excellent model for hair cell development and regeneration (Kniss et al., 2016; Nicolson, 2017; Pickett and Raible, 2019).

The lateral line system of zebrafish (a cyprinid teleost) is purely mechanosensory, as is the lateral line system of the other main anamniote lab model, the frog *Xenopus*. However, in many other vertebrates, the lateral line system also has an electrosensory division (Bullock et al., 1983; Baker et al., 2013; Crampton, 2019). In electroreceptive non-teleost jawed vertebrates, some or all of the neuromast lines on the head are flanked by fields of ampullary organs containing electroreceptors, which respond to weak cathodal stimuli such as the electric fields surrounding other animals in water (Bodznick and Montgomery, 2005; Crampton, 2019; Leitch and Julius, 2019; Chagnaud et al., 2021). Electroreception is mediated by voltage-gated calcium channels in the apical membrane (Bodznick and Montgomery, 2005; Leitch and Julius, 2019). The voltage sensor was recently identified in cartilaginous fishes as the L-type voltage-gated calcium channel Ca_v_1.3 (Bellono et al., 2017; Bellono et al., 2018), whose pore-forming alpha subunit is encoded by *Cacna1d*. Ca_v_1.3 is also required for synaptic transmission at hair-cell ribbon synapses (Moser et al., 2020; Mukhopadhyay and Pangrsic, 2022).

Electroreceptors, like hair cells, have an apical primary cilium and basolateral ribbon synapses with lateral line afferents (Jørgensen, 2005; Baker, 2019). However, in contrast to the highly ordered, stepped array of apical microvilli (’stereocilia’) that forms the ‘hair bundle’ critical for hair-cell mechanotransduction (Caprara and Peng, 2022), electroreceptors in many species (e.g., cartilaginous fishes; ray-finned paddlefishes and sturgeons) lack apical microvilli altogether (Jørgensen, 2005; Baker, 2019). Electroreceptors in other species have a few apical microvilli, while the electroreceptors of the amphibian axolotl have around 200 microvilli surrounding an eccentrically positioned primary cilium (Jørgensen, 2005; Baker, 2019). Indeed, axolotl electroreceptors were described as "remarkably similar to immature hair cells" (Northcutt et al., 1994). Thus, despite their shared function, the apical surface of electroreceptors (where voltage-sensing occurs; Bodznick and Montgomery, 2005; Leitch and Julius, 2019) varies considerably across different vertebrate groups.

Fate-mapping experiments have shown that neuromasts, ampullary organs (where present) and their afferent neurons all develop from a series of pre-otic and post-otic lateral line placodes on the embryonic head (Northcutt, 1997; Piotrowski and Baker, 2014; Baker, 2019). In electroreceptive jawed vertebrates, lateral line placodes elongate to form sensory ridges that eventually fragment: neuromasts differentiate first, in a line along the centre of each ridge, and ampullary organs (if present) form later, in fields on the flanks of the ridge (Northcutt, 1997; Piotrowski and Baker, 2014; Baker, 2019). The lateral line primordia of electroreceptive vertebrates therefore provide a fascinating model for studying the formation of different sensory cell types and organs. What molecular mechanisms control the formation within the same primordium of mechanosensory neuromasts containing hair cells, versus electrosensory ampullary organs containing electroreceptors?

To gain molecular insight into electroreceptor development, we originally took a candidate-gene approach, based on genes known to be important for neuromast and/or hair cell development. This enabled us to identify a variety of genes expressed in developing ampullary organs as well as neuromasts, in embryos from the three major jawed-vertebrate groups, i.e., cartilaginous fishes (lesser-spotted catshark, *Scyliorhinus canicula*, and little skate, *Leucoraja erinacea*; O’Neill et al., 2007; Gillis et al., 2012); lobe-finned bony fishes/tetrapods (a urodele amphibian, the axolotl, *Ambystoma mexicanum*; Modrell and Baker, 2012); and ray-finned bony fishes (a chondrostean, the Mississippi paddlefish, *Polyodon spathula*; Modrell et al., 2011a; Modrell et al., 2011b; Modrell et al., 2017b). We also took an unbiased discovery approach using differential bulk RNA-seq in late-larval paddlefish, which yielded a dataset of almost 500 genes that were putatively enriched in lateral line organs (Modrell et al., 2017a). Validation by *in situ* hybridization of a subset of candidates from this dataset suggested that conserved molecular mechanisms were involved in hair cell and electroreceptor development, and that hair cells and electroreceptors were closely related physiologically (Modrell et al., 2017a). For example, developing ampullary organs express the key ‘hair cell’ transcription factor genes *Atoh1* and *Pou4f3* (see Roccio et al., 2020; Iyer and Groves, 2021), and genes essential for the function of hair cell ribbon synapses, including the voltage-gated calcium channel gene *Cacna1d*, encoding Ca_v_1.3 (Modrell et al., 2017a). We also identified a handful of genes expressed in developing ampullary organs but not neuromasts, including two electroreceptor-specific voltage-gated potassium channel subunit genes (*Kcna5* and *Kcnab3*) and a single transcription factor gene, *Neurod4* (Modrell et al., 2017a).

Up to that point, we had reported the shared expression of fifteen transcription factor genes in both ampullary organs and neuromasts, but only one transcription factor gene with restricted expression, namely, electrosensory-restricted *Neurod4* (Modrell et al., 2011a; Modrell et al., 2011b; Modrell et al., 2017b). More recently (preprint: Minařík et al., 2023), we used the late-larval paddlefish lateral line organ-enriched dataset (Modrell et al., 2017b), as well as a candidate gene approach, to identify 23 more transcription factor genes expressed within developing lateral line organs in paddlefish and/or in a related, more experimentally tractable chondrostean, the sterlet (*Acipenser ruthenus*, a small sturgeon; e.g., Chen et al., 2018; Baloch et al., 2019; Stundl et al., 2022). Twelve of these transcription factor genes - including *Gfi1* - were expressed in both ampullary organs and neuromasts (preprint: Minařík et al., 2023). Thus, developing ampullary organs, as well as neuromasts, express the three ‘hair cell’ transcription factor genes - *Atoh1*, *Pou4f3* and *Gfi1* - whose co-expression is sufficient to drive postnatal mouse cochlear supporting cells to adopt a ‘hair cell-like’ fate, albeit not to form fully mature hair cells (Roccio et al., 2020; Iyer and Groves, 2021; Chen et al., 2021; Iyer et al., 2022). We also identified six novel ampullary organ-restricted transcription factor genes and the first-reported mechanosensory-restricted transcription factor genes (preprint: Minařík et al., 2023). One of the five mechanosensory-restricted transcription factor genes was *Foxg1*, which was expressed and maintained in the central region of sensory ridges where lines of neuromasts form, although excluded from hair cells (preprint: Minařík et al., 2023).

Here, we used CRISPR/Cas9-mediated mutagenesis in F0-injected sterlet embryos to investigate the function in lateral line organ development of *Atoh1*, electrosensory-restricted *Neurod4* and mechanosensory-restricted *Foxg1* (for reference, Supplementary Figure S1 shows the normal expression patterns of these genes). We report that *Atoh1* is required for the formation of electroreceptors, as well as hair cells. We did not see any phenotype after targeting ampullary organ-restricted *Neurod4*, potentially owing to redundancy with other *Neurod* family members that we found to be expressed in sterlet ampullary organs (and neuromasts). Targeting mechanosensory-restricted *Foxg1* resulted in a striking phenotype: the ectopic formation of ampullary organs within neuromast lines, and in some cases the fusion of ampullary organ fields that normally develop either side of a line of neuromasts. This suggests the unexpected but intriguing hypothesis that ampullary organs may be the ‘default’ fate for lateral line sensory ridges, and that this is repressed by Foxg1, allowing neuromasts to form instead.

## Results

### CRISPR/Cas9-mediated mutagenesis in F0-injected sterlet embryos

To test gene function during lateral line organ development, we optimised CRISPR/Cas9-mediated mutagenesis in F0-injected sterlet embryos, building on established protocols for axolotl (*Ambystoma mexicanum*; Flowers et al., 2014; Fei et al., 2018), newt (*Pleurodeles waltl*; Elewa et al., 2017), and sea lamprey (*Petromyzon marinus*; Square et al., 2015; York et al., 2019; Square et al., 2020), whose eggs are all large (1-2 mm in diameter) and easy to microinject at the 1-2 cell stage. Ovulated sterlet eggs are very large: roughly 2.5 mm in diameter (Lenhardt et al., 2005). (Since this project started, CRISPR/Cas9-mediated mutagenesis in F0-injected sterlet embryos has been reported, including by two of us, R.F. and M.P.; Chen et al., 2018; Baloch et al., 2019; Stundl et al., 2022.) Analysis of microsatellite data had originally suggested that although a whole-genome duplication had occurred in the sterlet lineage, the sterlet was likely to be a functional diploid (Ludwig et al., 2001). Our sgRNAs were designed before the first chromosome-level sterlet genome was published (Du et al., 2020). Analysis of this genome showed that approximately 70% of ohnologs (i.e., gene paralogs originating from the whole-genome duplication) had in fact been retained, suggesting functional tetraploidy (Du et al., 2020). We comment on this in relation to our experiments at the relevant points below.

We targeted the melanin-producing enzyme *tyrosinase* (*Tyr*) as a positive control, using eight different single-guide (sg) RNAs. Table 1 shows the sgRNA target sequences (including two that were designed and recently published by Stundl et al., 2022) and the various combinations in which they were injected. Supplementary Figure S2A shows the position of each sgRNA relative to the exon structure of the *tyrosinase* gene. Our sgRNAs were designed before chromosome-level sterlet genomes were available (Du et al., 2020 and the 2022 reference genome: https://www.ncbi.nlm.nih.gov/datasets/genome/GCF_902713425.1). Searching the reference genome for *Tyr* showed that both *Tyr* ohnologs have been retained, on chromosomes 8 and 9, with 99.11% nucleotide identity in the coding sequence (98.81% amino acid identity). Our sgRNAs target both *Tyr* ohnologs equally.

**Table 1.**
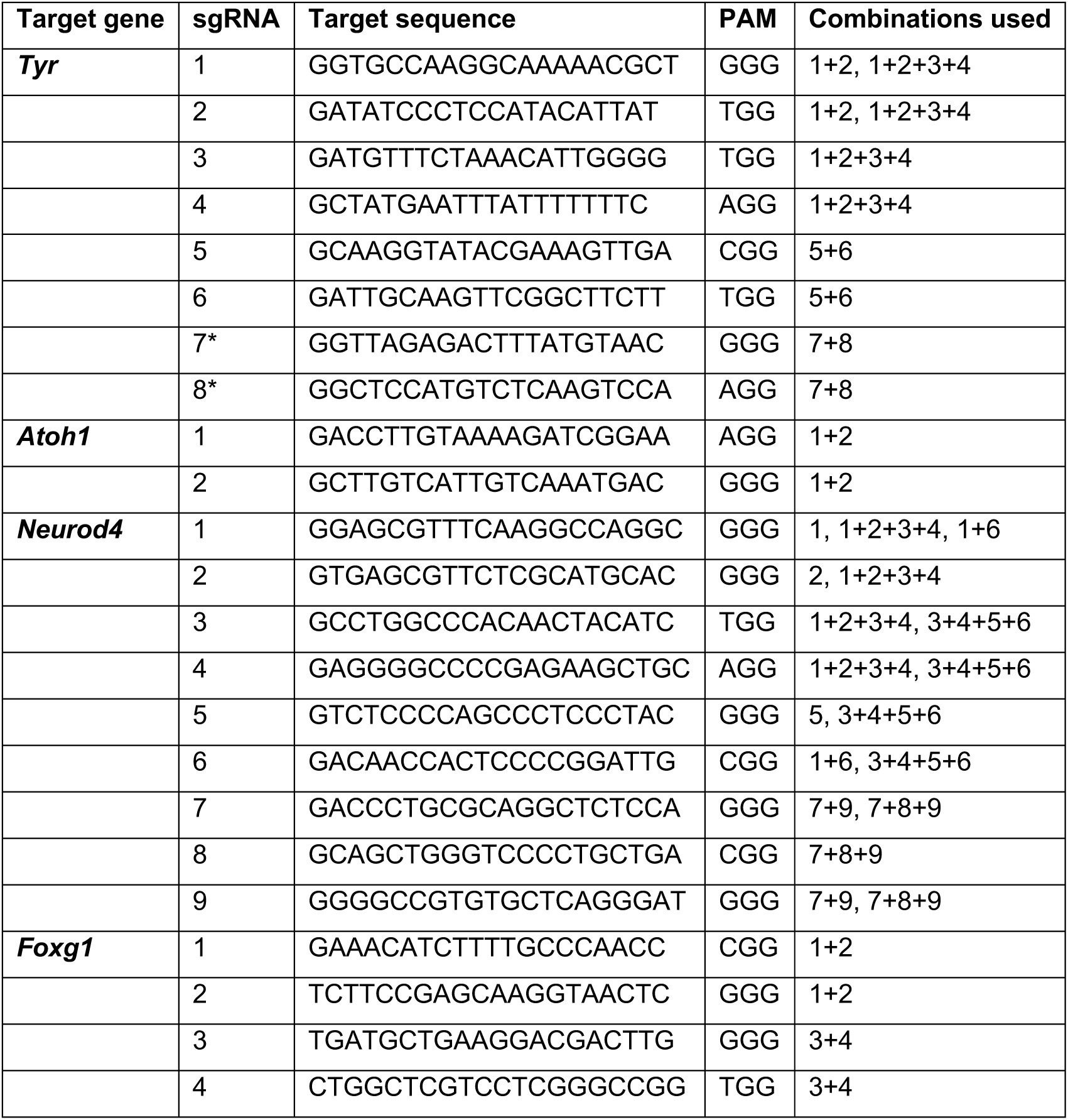
sgRNAs used in this study. List of the genes targeted for CRISPR/Cas9-mediated mutagenesis, together with the target sequences and combinations of sgRNAs reported in this study. *Note: Tyr* sgRNAs 7 and 8 (marked with an asterisk) were designed and recently published by Stundl et al. (2022) as their *tyr* sgRNA 3 and *tyr* sgRNA 4, respectively.

Three different combinations of six of the eight *Tyr* sgRNAs, when injected at the 1-2- cell stage (2-4 sgRNAs pre-complexed with Cas9), each generated at least four embryos (hereafter ‘crispants’) with altered pigmentation phenotypes evaluated at stage 45 (Dettlaff et al., 1993), the onset of the transition to independent feeding. The other two *Tyr* sgRNAs failed to generate any phenotypes (Supplementary Table 1). Excluding the *Tyr* sgRNAs that failed, at least some degree of pigment loss was seen in 63/111 *Tyr* crispants (56.8%) across nine independent batches. Examples of *Tyr* crispants with pigmentation phenotypes, plus a control, are shown in Supplementary Figure S2B-D. The most efficient results were obtained by injecting 1-cell embryos with a preassembled mix of Cas9 protein plus two chemically modified sgRNAs (purchased from Synthego) against the target gene, and subsequently maintaining the embryos at room temperature for around 6 hours. The time from fertilization to completion of the first cleavage is around 2-3 hours at room temperature, giving plenty of time for the Cas9/sgRNA complex to act before returning the embryos to colder temperatures for subsequent development. (Sterlet are cold-water fish and the optimum temperature for maintaining embryos for normal development is 16 °C.)

Following embryo injection at the 1-2-cell stage with pre-complexed sgRNAs/Cas9 and fixation at stage 45, genomic DNA was extracted from the trunk/tail prior to analysis of the heads by *in situ* hybridization (ISH). The sgRNA-targeted region from trunk/tail genomic DNA was amplified by PCR for direct Sanger sequencing and *in silico* analysis using Synthego’s online ‘Inference of CRISPR Edits’ (ICE) tool (Conant et al., 2022) (also see e.g., Uribe-Salazar et al., 2022) to analyse the identity and frequency of edits of the target gene. Although our genotyping primers were designed before chromosome-level sterlet genomes were available, comparison with the reference genome showed no mismatches against either of the two ohnologs. Genotyping and ICE analysis (Conant et al., 2022) of tails from individual *Tyr* crispants showed successful disruption of the *Tyr* gene (Supplementary Figure S2E-I show examples of successful disruption of *Tyr*; the genotyping data were consistent with the primers amplifying both ohnologs).

We note that our genotyping results have shown that most crispants analysed, across all genes targeted, have shown some degree of targeted mutagenesis in the trunk/tail, with a range of deletion sizes. Although phenotypes from the initial spawning seasons were almost always highly mosaic, suggesting mutations occurred later in development, following optimization a proportion of embryos showed complete unilateral and occasionally bilateral phenotypes. Such phenotypes suggest that mutation occurred in one cell at the 2-cell stage (unilateral phenotype) or even as early as the 1-cell stage (bilateral phenotype). Some degree of mosaicism can be useful, however, as the unaffected tissue provides an internal control for the normal expression of the gene being examined by ISH.

### Targeting *Atoh1* resulted in the loss of hair cells and electroreceptors

We targeted *Atoh1* for CRISPR/Cas9-mediated mutagenesis by injecting sterlet embryos at the 1-2 cell stage with Cas9 protein pre-complexed with two sgRNAs targeting *Atoh1* (Table 1; Supplementary Figure S2A). Our sgRNAs were designed before chromosome-level sterlet genomes were available (Du et al., 2020 and the 2022 reference genome: https://www.ncbi.nlm.nih.gov/datasets/genome/GCF_902713425.1/). Searching the reference genome for *Atoh1* showed that both *Atoh1* ohnologs have been retained, on chromosomes 1 and 2, with 91.41% nucleotide identity (and 84.02% amino acid identity) in the coding sequence. The copy on chromosome 2 encodes a shorter version of the protein with a four amino acid deletion near the N-terminus (E14_G17del). Our *Atoh1* sgRNA 1 (Table 1; Supplementary Figure S3A) has a one-base mismatch to the shorter *Atoh1* gene on chromosome 2, in position 3 of the target sequence (PAM-distal), which is unlikely to prevent successful targeting (Wu et al., 2014). Our *Atoh1* sgRNA 2 (Table 1; Supplementary Figure S3A) has a two-base mismatch to the longer *Atoh1* gene on chromosome 1, in positions 1 and 2 of the target sequence (PAM-distal), which is also unlikely to prevent successful targeting. Thus, we expect our sgRNAs to target both *Atoh1* ohnologs.

The *Atoh1* crispants were raised to stage 45 (the onset of independent feeding, around 14 days post-fertilization), together with *Tyr*-targeted siblings/half-siblings as controls (eggs were fertilized *in vitro* with a mix of sperm from three different males). ISH for the hair cell and electroreceptor marker *Cacna1d* (Modrell et al., 2017a) (also see preprint: Minařík et al., 2023), which was recently shown to be a direct Atoh1 target gene in mouse cochlear hair cells (Jen et al., 2022), revealed no obvious lateral line organ phenotype in *Tyr* control crispants (Figure 1A-D; n=0/17 across four independent batches; Supplementary Table 1). Even in wildtype larvae, the number of ampullary organs in individual fields varies considerably at stage 45, so ampullary organ number was not in itself scored as a phenotype. However, *Cacna1d* expression was absent mosaically in neuromast lines and ampullary organ fields in *Atoh1* crispants (Figure 1E-J; n=13/22, i.e., 59%, across five independent batches; Supplementary Table 1). This suggested that disruption of the *Atoh1* gene in sterlet resulted in the failure of hair cell differentiation (as expected from zebrafish; Millimaki et al., 2007) and also electroreceptors. Post-ISH immunostaining for the supporting-cell marker Sox2 (Hernández et al., 2007; Modrell et al., 2017a) (also see preprint: Minařík et al., 2023) confirmed that neuromasts had formed (Figure 1E^1^,F^1^), so the phenotype was specific to receptor cells. Ampullary organs were not reliably detected because Sox2 immunostaining labels neuromasts much more strongly than ampullary organs in sterlet (Supplementary Figure S1A,B; also preprint: Minařík et al., 2023), as it does in paddlefish (Modrell et al., 2017a). ISH for electroreceptor-specific *Kcnab3* (Modrell et al., 2017a) (also see preprint: Minařík et al., 2023) similarly showed no effect in *Tyr* control crispants (Figure 1K-N; n=0/27 across six independent batches; Supplementary Table 1), but the mosaic absence of *Kcnab3* expression in ampullary organ fields in *Atoh1* crispants (Figure 1O-T; n=8/13, i.e., 62%, across two independent batches; Supplementary Table 1). Post-*Kcnab3* ISH immunostaining for Sox2 confirmed that ampullary organs were still present, as well as neuromasts (Figure 1O^1^,P^1^).

**Figure 1.**
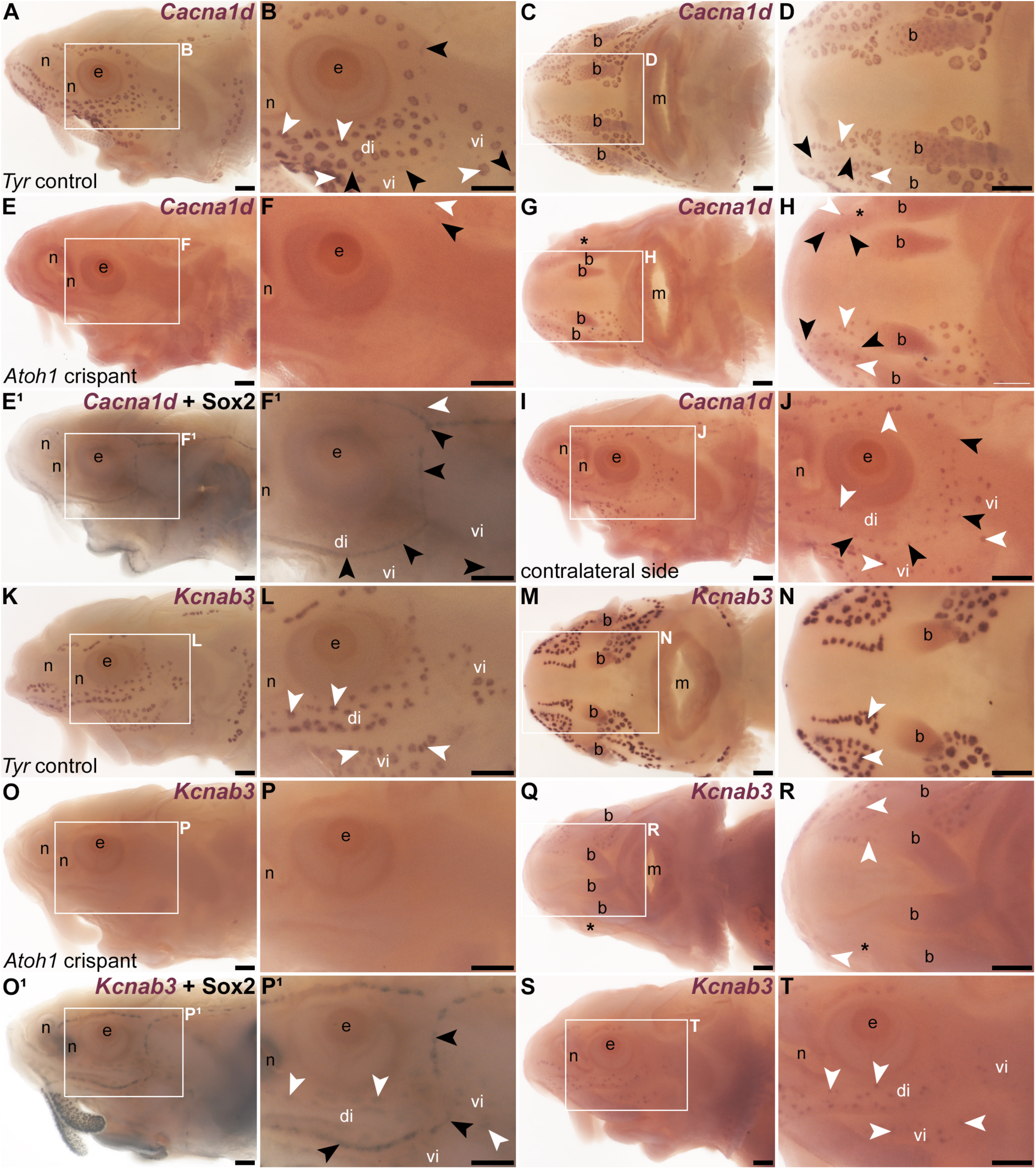
*Atoh1* is required for the differentiation of lateral line hair cells and electroreceptors. Sterlet crispants at stage 45 after *in situ* hybridization (ISH) for the hair cell and electroreceptor marker *Cacna1d*, or the electroreceptor-specific marker *Kcnab3*. Black arrowheads indicate examples of neuromasts; white arrowheads indicate examples of ampullary organs. (**A-D**) In a control *Tyr* crispant, *Cacna1d* expression shows the normal distribution of hair cells in lines of neuromasts, and electroreceptors in fields of ampullary organs flanking the neuromast lines (lateral view: A,B; ventral view: C,D). (**E-J**) In an *Atoh1* crispant (from a different batch to the *Tyr* crispant shown in A-D), *Cacna1d* expression is absent on the left side of the head (E,F), except for a few isolated organs in the otic region and on the operculum. Post-ISH Sox2 immunostaining (E^1^,F^1^) shows that neuromast supporting cells are still present. A ventral view (G,H) and a lateral view of the right side of the head (I,J; image flipped horizontally for ease of comparison) reveal a unilateral phenotype, with *Cacna1d-*expressing hair cells and electroreceptors mostly absent from the left side only of the ventral rostrum (asterisk in G,H) and present on the right side of the head (I,J). (**K,L**) Lateral view of a control *Tyr* crispant after ISH for *Kcnab3*, showing the position of electroreceptors in ampullary organs. (**M,N**) Ventral view of another *Tyr* crispant showing *Kcnab3* expression in ampullary organs. (**O-P^1^**) Lateral view of an *Atoh1* crispant in which *Kcnab3* expression is absent from ampullary organs. Post-ISH Sox2 immunostaining (O^1^,P^1^) shows that supporting cells are still present in neuromasts (strong staining) and can also be detected in ampullary organs (much weaker staining). (**Q-T**) A different *Atoh1* crispant after ISH for *Kcnab3*, shown in ventral view (Q,R: compare with M,N) and lateral view (S,T). *Kcnab3* expression reveals a unilateral phenotype: *Kcnab3-*expressing electroreceptors are mostly absent from the right side (asterisk) but present on the left side. Abbreviations: b, barbel; di, dorsal infraorbital ampullary organ field; e, eye; m, mouth; n, naris; S, stage; vi, ventral infraorbital ampullary organ field. Scale bars: 200 μm.

In mouse cochlear hair cells, the ‘hair cell’ transcription factor genes *Pou4f3* and *Gfi1* are direct Atoh1 targets (Yu et al., 2021; Jen et al., 2022). ISH for *Pou4f3* showed no phenotype in *Tyr* controls (Figure 2A,B; n=0/9 across three batches; Supplementary Table 1) but the mosaic absence of *Pou4f3* in ampullary organ fields and neuromast lines of *Atoh1* crispants (Figure 2C-D^1^; n=13/15 embryos, i.e., 87%, across three batches; Supplementary Table 1). Similarly, ISH for *Gfi1* showed no phenotype in *Tyr* controls (Figure 2E,F; n=0/17 across five batches; Supplementary Table 1), but mosaic absence of *Gfi1* in ampullary organ fields and neuromast lines of *Atoh1* crispants (Figure 2G-H^1^; n=9/14 embryos, i.e., 64%, across four batches; Supplementary Table 1). Post-ISH immunostaining for Sox2 confirmed that neuromasts and ampullary organs were still present in *Atoh1* crispants (Figure 2C^1^,D^1^,G^1^,H^1^).

**Figure 2.**
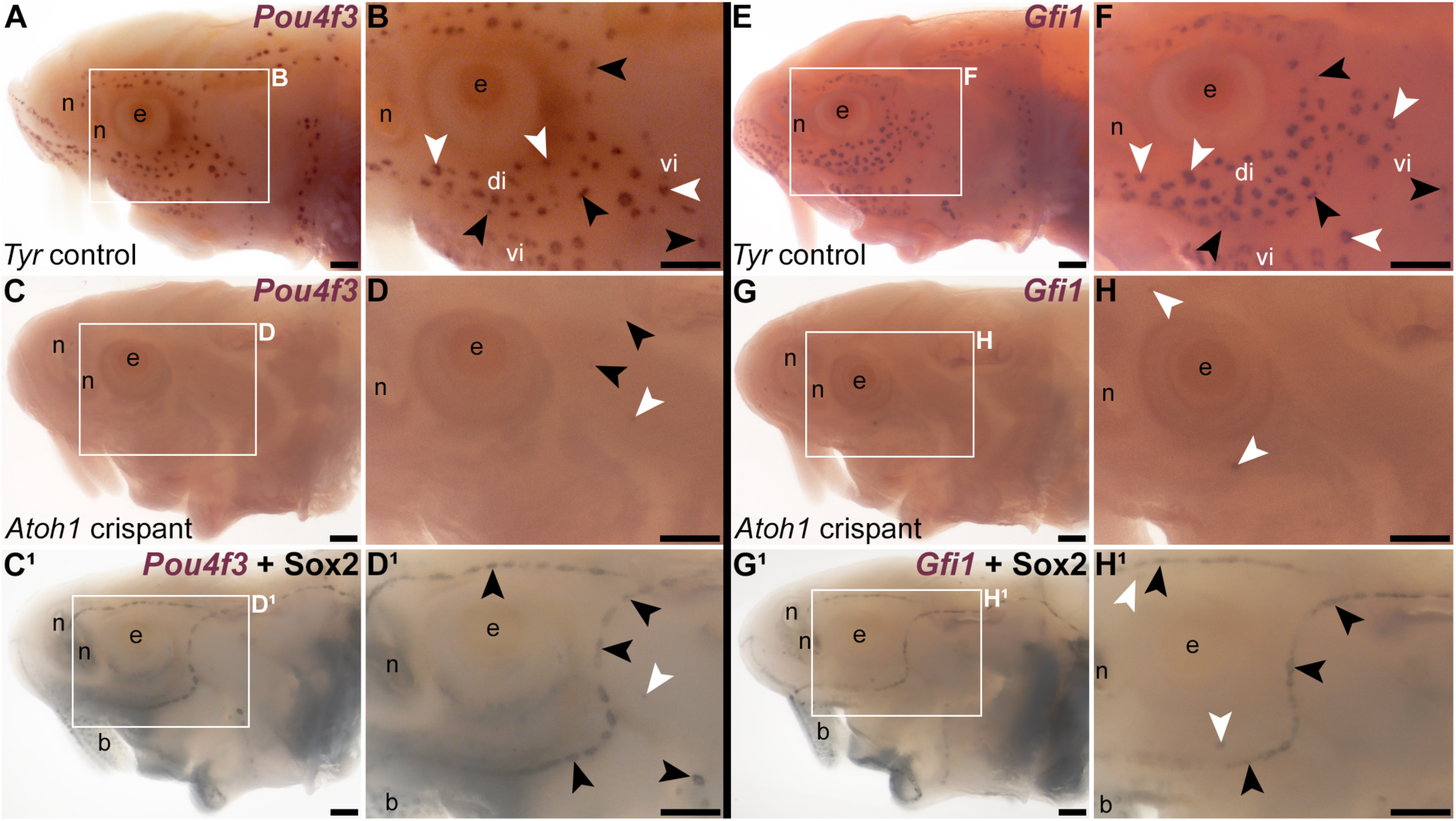
*Atoh1* is required for *Pou4f3* and *Gfi1* expression in ampullary organs and neuromasts. Sterlet crispants at stage 45 after *in situ* hybridization for transcription factor genes expressed by developing hair cells. Black arrowheads indicate examples of neuromasts; white arrowheads indicate examples of ampullary organs. (**A,B**) In a control *Tyr* crispant, *Pou4f3* expression is detected in both neuromasts and ampullary organs. (**C-D^1^**) In an *Atoh1* crispant, *Pou4f3* expression is absent from both neuromasts and ampullary organs, except for a few isolated organs in the postorbital region. Post-ISH Sox2 immunostaining (C^1^,D^1^) shows that neuromast supporting cells are still present. Most ampullary organs are not visible as Sox2 expression in ampullary organs is significantly weaker than in neuromasts (Supplementary Figure S1A,B) and often not detectable after post-ISH immunostaining. (**E,F**) In a control *Tyr* crispant, *Gfi1* expression is detected in both neuromasts and ampullary organs. (**G-H^1^**) In an *Atoh1* crispant, *Gfi1* expression is absent from both neuromasts and ampullary organs, except for a few isolated organs in the supra- and infraorbital region. Post-ISH Sox2 immunostaining (G^1^,H^1^) shows that neuromast supporting cells are still present. (Most ampullary organs are not visible due to weaker Sox2 immunostaining.) Abbreviations: b, barbel; di, dorsal infraorbital ampullary organ field; e, eye; n, naris; S, stage; vi, ventral infraorbital ampullary organ field. Scale bars: 200 μm.

We performed genotyping and ICE analysis (Conant et al., 2022) of tails from individual *Atoh1* crispants targeted with this pair of sgRNAs (Supplementary Figure S3 shows examples). Our genotyping primers were designed before chromosome-level sterlet genomes were available; comparison with the reference genome showed two mismatches in the forward primer against the ohnolog on chromosome 2, and the genotyping data were consistent with the primers amplifying the chromosome 1 ohnolog only. Thus, we could not determine whether the chromosome 2 ohnolog was disrupted. However, the genotyping data showed successful disruption of the *Atoh1* gene on chromosome 1 (Supplementary Figure S3).

Taken together, these data suggest that Atoh1 lies upstream of *Pou4f3* and *Gfi1* in ampullary organs as well as neuromasts, and is required for the differentiation of electroreceptors as well as hair cells.

### Targeting electrosensory-restricted *Neurod4* had no obvious effect on lateral line development

We previously identified *Neurod4* in paddlefish as the first-reported ampullary organ-restricted transcription factor in the developing lateral line system (Modrell et al., 2017a). We confirmed that sterlet *Neurod4* is similarly expressed by ampullary organs but not neuromasts (Supplementary Figure S1I,J). We targeted *Neurod4* in sterlet embryos using nine different sgRNAs (Table 1; Supplementary Figure S4A), injected in eight different combinations of 1-4 different sgRNAs across 10 independent batches of 1-2 cell-stage embryos (Supplementary Table 1). This had no detectable effect on expression of electroreceptor-specific *Kcnab3* (n=0/44 across nine batches, Supplementary Table 1) or the hair cell/electroreceptor marker *Cacna1d* (n=0/4 across two batches, Supplementary Table 1). Examples of *Neurod4* crispants after ISH for *Kcnab3*, plus a *Tyr* control crispant for comparison, are shown in Supplementary Figure S4B-D. Our sgRNAs were designed before chromosome-level sterlet genomes were available (Du et al., 2020 and the 2022 reference genome: https://www.ncbi.nlm.nih.gov/datasets/genome/GCF_902713425.1/). Searching the reference genome for *Neurod4* showed that both ohnologs have been retained: one on chromosome 45 and the other annotated on an unplaced genomic scaffold, with 99.45% nucleotide identity (and 99.46% amino acid identity) in the coding sequence. Our sgRNAs target both *Neurod4* ohnologues without mismatches. Our genotyping primers were designed before chromosome-level sterlet genomes were available; comparison with the reference genome showed five mismatches in the forward primer used for genotyping larvae targeted with sgRNAs 1 and 2 against the chromosome 45 ohnolog (Supplementary Table 1). It was not possible to tell from our genotyping data whether the primers amplified both ohnologs or only one, however, as the remaining sequence targeted by the primers is identical between the two ohnologs. Genotyping and ICE analysis (Conant et al., 2022) showed successful disruption of the *Neurod4* gene arising from six different combinations of the sgRNAs (Supplementary Table 1; examples are shown in Supplementary Figure S4E-I). Not all sgRNA combinations were genotyped, but all *Neurod4* crispants counted (n=44) included at least one sgRNA confirmed to disrupt the *Neurod4* gene via genotyping of other embryos (Supplementary Table 1).

The lack of phenotype in *Neurod4* crispants, despite successful disruption of the *Neurod4* gene, suggested either that Neurod4 is not required for electroreceptor differentiation, despite its restriction to ampullary organs in both paddlefish and sterlet (Modrell et al., 2017a; this paper), or that it acts redundantly with other transcription factors. In paddlefish, *Neurod1* expression was restricted to developing lateral line ganglia (Modrell et al., 2011b). We cloned sterlet *Neurod1*, *Neurod2* and *Neurod6*. (Unlike *Neurod4*, these three *Neurod* family members are all direct Atoh1 targets in mouse hair cells; Jen et al., 2022.) These three genes all proved to be expressed in sterlet ampullary organs, as well as neuromasts (Supplementary Figure S5). Thus, it seems likely that the lack of effect of CRISPR/Cas9-mediated targeting of sterlet *Neurod4* is due to redundancy with other Neurod family transcription factors co-expressed in ampullary organs. (Our results also show there is at least some variation in *Neurod* family gene expression within the developing lateral line systems of paddlefish and sterlet.)

### Targeting mechanosensory-restricted *Foxg1* led to the formation of ectopic ampullary organs within neuromast lines

We recently identified *Foxg1* as a mechanosensory lateral line-restricted transcription factor gene in paddlefish and sterlet (preprint: Minařík et al., 2023). *Foxg1* is expressed in the central zones of lateral line sensory ridges where neuromasts form, though excluded from hair cells themselves (preprint: Minařík et al., 2023). Our sgRNAs against *Foxg1* (Table 1; Supplementary Figure S6A) were designed before chromosome-level sterlet genomes were available (Du et al., 2020 and the 2022 reference genome (https://www.ncbi.nlm.nih.gov/datasets/genome/GCF_902713425.1/). Searching the reference genome for *Foxg1* showed that both *Foxg1* ohnologs have been retained, on chromosomes 15 and 18, with 96.43% nucleotide identity in the coding sequence (99.51% amino acid identity). Our sgRNAs target both *Foxg1* ohnologs without mismatches.

When compared with *Tyr* control crispants (Figure 3A-D), a striking phenotype was seen mosaically after targeting mechanosensory-restricted *Foxg1* with sgRNAs 1 and 2 (Table 1; Supplementary Figure S6A): neuromast lines were often interrupted by ectopic ampullary organs. These were defined as *Cacna1d*-expressing cells present within neuromast lines in larger clusters than expected for neuromasts, resembling ampullary organs (Figure 3E-L; n=9/18, i.e., 50%, across two independent batches; Supplementary Table 1), or by expression of electroreceptor-specific *Kcnab3* within neuromast lines, not seen in *Tyr* control crispants (Figure 3M-V; n=7/24, i.e., 29%, across both batches; Supplementary Table 1). This sometimes led to the apparent fusion of ampullary organ fields across a missing neuromast line (e.g., Figure 3E-H,S-V). Post-*Kcnab3* ISH immunostaining for Sox2, to reveal neuromasts, also confirmed that the ectopic electroreceptors in *Foxg1* crispants were within neuromast lines (Figure 3Q^1^,R^1^). Targeting *Foxg1* with a different pair of sgRNAs (Table 1; Supplementary Figure S6A) in a different batch of embryos generated similar phenotypes in 3/21 embryos (14%) overall (*Cacna1d*: n=1/6, i.e., 17%; *Kcnab3*: n=2/15, i.e., 13%; Supplementary Table 1).

**Figure 3.**
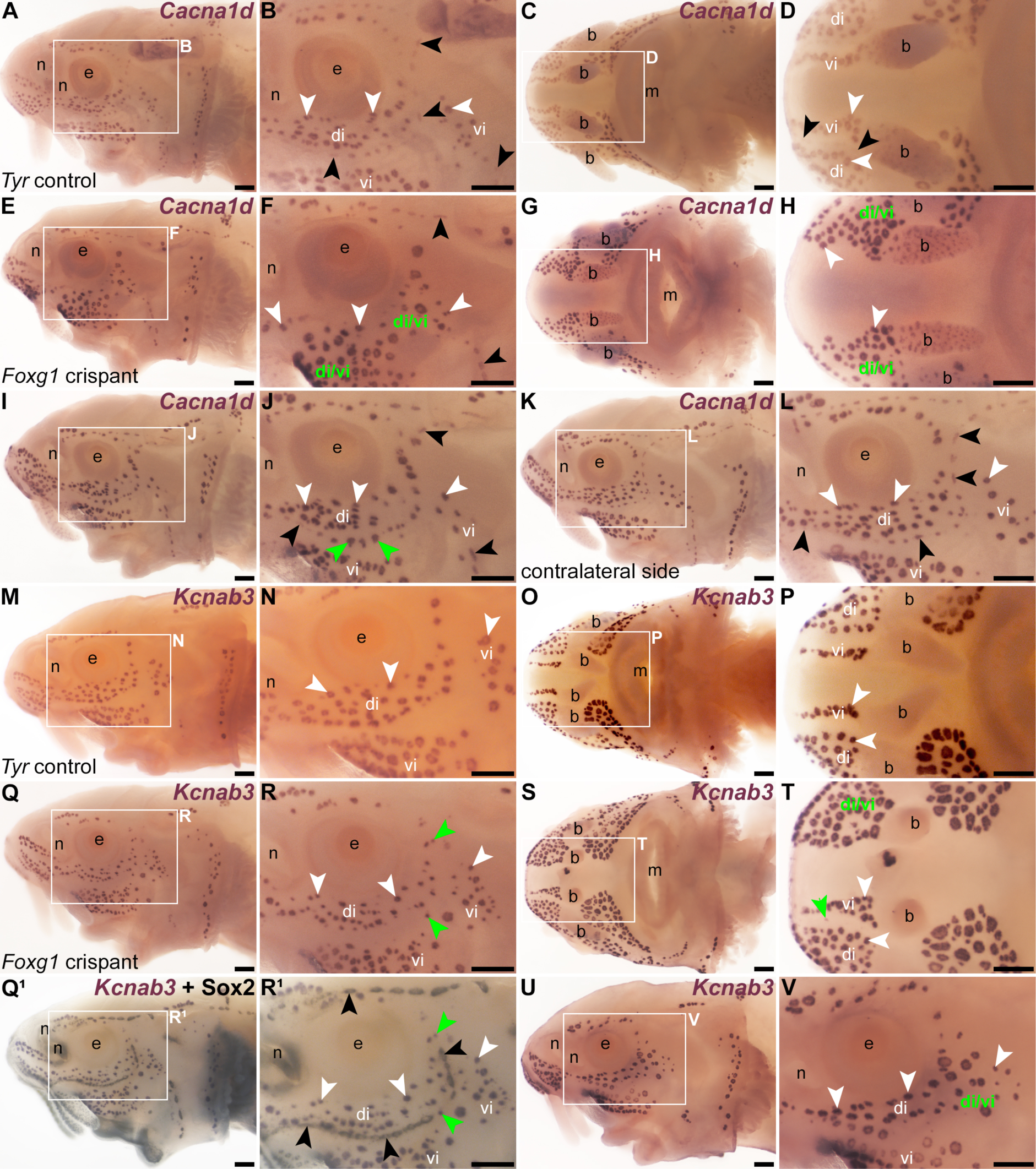
*Foxg1* represses ampullary organ and electroreceptor formation within neuromast lines. Sterlet crispants at stage 45 after *in situ* hybridization (ISH) for the hair cell and electroreceptor marker *Cacna1d*, or the electroreceptor-specific marker *Kcnab3*. Black arrowheads indicate examples of neuromasts; white arrowheads indicate examples of ampullary organs. (**A,B**) Lateral view of a control *Tyr* crispant. *Cacna1d* expression shows the normal distribution of hair cells and electroreceptors. Note that ampullary organs have significantly more *Cacna1d*-expressing receptor cells than neuromasts. (**C,D**) Ventral view of a second control *Tyr* crispant. *Cacna1d* expression reveals the infraorbital neuromast line on both sides of the ventral rostrum, flanked by the dorsal infraorbital (di) and ventral infraorbital (vi) ampullary organ fields. (**E,F**) Lateral view of a *Foxg1* crispant. *Cacna1d* expression reveals that distinct neuromast lines are missing and the corresponding space is filled by putative ectopic ampullary organs, based on the large, ampullary organ-like clusters of *Cacna1d*-expressing cells. The dorsal and ventral infraorbital ampullary organ fields seem to have fused across the missing neuromast line (compare with A,B). (**G,H**) Ventral view of a second *Foxg1* crispant. *Cacna1d* expression reveals an apparent fusion of the dorsal infraorbital (di) and ventral infraorbital (vi) ampullary organ fields across the missing infraorbital neuromast lines on both sides (compare with C,D). (**I-K**) In a third *Foxg1* crispant, *Cacna1d* expression on the left side (I,J) shows that distinct supraorbital and infraorbital neuromast lines are still present. However, some organs within the supraorbital line and most organs within the infraorbital line have large clusters of *Cacna1d*-expressing cells, suggesting they are ectopic ampullary organs (green arrowheads in J show examples). On the right side (K,L; image flipped horizontally for ease of comparison), this phenotype is not seen. (**M,N**) Lateral view of a third control *Tyr* crispant. Electroreceptor-specific *Kcnab3* expression shows the distribution of ampullary organs only. (**O,P**) Ventral view of a fourth control *Tyr* crispant. *Kcnab3* expression shows the distribution of ampullary organ fields. Note the lack of staining where the infraorbital neuromast lines run on either side of the ventral rostrum, flanked by the dorsal infraorbital (di) and ventral infraorbital (vi) ampullary organ fields (compare with *Cacna1d* expression in C,D). (**Q-R^1^**) Lateral view of a fourth *Foxg1* crispant. *Kcnab3* expression shows two ectopic ampullary organs (green arrowheads) within the infraorbital neuromast line (compare with M,N). Post-ISH Sox2 immunostaining for supporting cells (Q^1^,R^1^) confirms the position of the neuromast lines. (**S,T**) Ventral view of a fifth *Foxg1* crispant. On the left side, ectopic *Kcnab3-*expressing electroreceptors fill the space where the left infraorbital neuromast line would normally run, such that the dorsal and ventral infraorbital ampullary organ fields seem to have fused (compare with O,P). On the right side, a single ectopic *Kcnab3-*expressing ampullary organ is visible (green arrowhead) where the right infraorbital neuromast line runs (compare with O,P). (**U,V**) Lateral view of a sixth *Foxg1* crispant. *Kcnab3* expression shows fusion of the dorsal infraorbital ampullary organ field with the dorsal part of the ventral infraorbital ampullary organ field (compare with M,N). Abbreviations: b, barbel; di, dorsal infraorbital ampullary organ field; di/vi, fused dorsal infraorbital and ventral infraorbital ampullary organ fields; e, eye; m, mouth; n, naris; S, stage; vi, ventral infraorbital ampullary organ field. Scale bars: 200 μm.

We also investigated this phenotype by performing ISH for two ampullary organ-restricted transcription factor genes, *Mafa* (preprint: Minařík et al., 2023) and *Neurod4* (Modrell et al., 2017a; this study), followed by Sox2 immunostaining in some embryos to reveal the location of neuromasts. Relative to the normal ampullary organ expression of *Mafa* in uninjected siblings/half-siblings (sufficient *Tyr* control crispants were not available to test; Figure 4A-D; n=0/6 within one batch), ISH for *Mafa* showed the mosaic presence of ampullary organs in neuromast lines and/or merging of ampullary organ fields in *Foxg1* crispants (Figure 4E-J; n=3/17 embryos i.e., 18%, across two independent batches; Supplementary Table 1). Similarly, relative to the usual ampullary organ expression of *Neurod4* in uninjected siblings/half-siblings (sufficient *Tyr* control crispants were not available to test; Figure 4K-N; n=0/2 embryos within one batch; Supplementary Table 1), ISH for *Neurod4* (more weakly expressed than *Mafa*) revealed the same ectopic ampullary organ phenotype in *Foxg1* crispants as seen for *Cacna1d*, *Kcnab3* and *Mafa* (Figure 4O-T; n=3/6 embryos, i.e., 50%, within one batch; Supplementary Table 1). Although our genotyping primers were designed before chromosome-level sterlet genomes were available, comparison with the reference genome showed no mismatches against either of the two ohnologs. Genotyping and ICE analysis (Conant et al., 2022) of tails from individual *Foxg1* crispants showed successful disruption of the *Foxg1* gene (Supplementary Figure S6 shows examples; the genotyping data were consistent with the primers amplifying both ohnologs). Overall, these data suggest that mechanosensory-restricted Foxg1 acts to repress the formation of ampullary organs and electroreceptors within neuromast lines.

**Figure 4.**
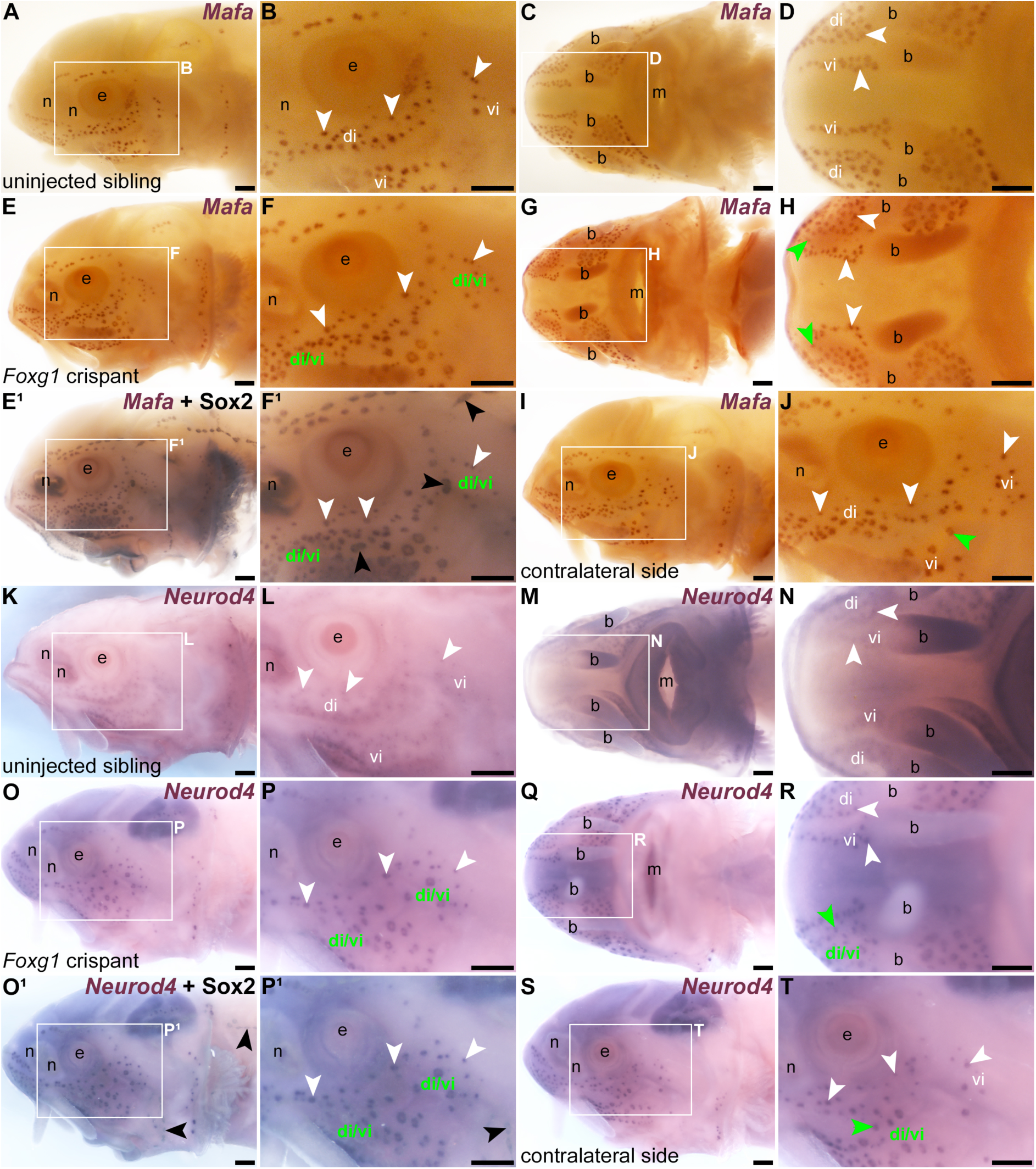
Ectopic ampullary organs in *Foxg1* crispants express ampullary organ-specific transcription factor genes *Mafa* and *Neurod4*. Sterlet crispants at stage 45 after *in situ* hybridization (ISH) for ampullary organ-restricted transcription factor genes. Black arrowheads indicate examples of neuromasts; white arrowheads indicate examples of ampullary organs. (**A-D**) In an uninjected sibling/half-sibling (eggs were fertilized *in vitro* with a mix of sperm from three different males), *Mafa* expression is restricted to ampullary organs (lateral view: A,B; ventral view: C,D). (**E-J**) A *Foxg1* crispant. On the left side of the head (E,F), several *Mafa*-expressing ectopic ampullary organs are present in the space where the infraorbital neuromast line would normally run, such that the dorsal and ventral infraorbital ampullary organ fields seem to have fused (compare with A,B). Post-ISH Sox2 immunostaining (E^1^,F^1^) shows that neuromasts are still present both proximally and distally to the sites of ampullary organ field fusion. In ventral view (G,H), ectopic ampullary organs (green arrowheads) are seen bilaterally, within the spaces where the infraorbital neuromast lines run on either side of the ventral rostrum (compare with C,D). On the right side in lateral view (I,J; image flipped horizontally for ease of comparison), a single *Mafa*-expressing ectopic ampullary organ (green arrowhead) is also present in the space where the infraorbital neuromast line runs (compare with A,B). (**K-N**) In an uninjected sibling/half-sibling, *Neurod4* expression is restricted to ampullary organs (lateral view: K,L; ventral view: M,N). (**O-T**) A *Foxg1* crispant. On the left side of the head (O,P), *Neurod4*-expressing ectopic ampullary organs are present in the space where the infraorbital neuromast line would normally run, such that the dorsal and ventral infraorbital ampullary organ fields seem to have fused (compare with K,L). Post-ISH Sox2 immunostaining (O^1^,P^1^) suggests that neuromasts are absent from the site of infraorbital ampullary organ field fusion, although neuromasts can be seen in the preopercular and trunk lines (black arrowheads, compare with O,P). In ventral view (Q,R), ectopic *Neurod4-*expressing ampullary organs are seen where the right infraorbital neuromast line would normally run on the ventral rostrum (green arrowhead indicates an example), resulting in partial fusion of the dorsal and ventral infraorbital fields on this side (the left side is unaffected). On the right side in lateral view (S,T; image flipped horizontally for ease of comparison), ectopic ampullary organs are also present in the space where the infraorbital neuromast line runs (green arrowhead in T indicates an example), resulting in the apparent partial fusion of the dorsal and ventral infraorbital ampullary organ fields (compare with K,L). Abbreviations: b, barbel; di, dorsal infraorbital ampullary organ field; di/vi, fused dorsal infraorbital and ventral infraorbital ampullary organ fields; e, eye; m, mouth; n, naris; S, stage; vi, ventral infraorbital ampullary organ field. Scale bars: 200 μm.

## Discussion

### Conserved molecular mechanisms underlie lateral line electroreceptor and hair cell formation

Here, we aimed to test the function in lateral line electroreceptor and/or hair cell formation of three transcription factor genes that we had previously identified as expressed in developing electrosensory ampullary organ and/or mechanosensory lateral line organs in ray-finned chondrostean fishes - paddlefish and sterlet (Butts et al., 2014; Modrell et al., 2017a; preprint: Minařík et al., 2023). The first gene we investigated was *Atoh1*, which is required for the formation of lateral line hair cells in zebrafish (Millimaki et al., 2007), as well as hair cells in the inner ear (Bermingham et al., 1999; Millimaki et al., 2007). In paddlefish (Butts et al., 2014; Modrell et al., 2017a) and sterlet (preprint: Minařík et al., 2023), *Atoh1* is expressed in ampullary organs as well as neuromasts. Targeting *Atoh1* for CRISPR/Cas9-mediated mutagenesis in F0 sterlet embryos showed that Atoh1 is required for the formation not only of *Cacna1d*-expressing neuromast hair cells, as expected from zebrafish (Millimaki et al., 2007), but also of *Cacna1d*-expressing, *Kcnab3*-expressing electroreceptors. These experiments also showed that Atoh1 is required for the expression of the ‘hair cell’ transcription factor genes *Gfi1* and *Pou4f3* in developing ampullary organs, as well as neuromasts. This is consistent with both of these genes being direct Atoh1 targets in mouse cochlear hair cells (Yu et al., 2021; Jen et al., 2022).

In both inner-ear hair cells and Merkel cells (epidermal mechanoreceptors found in all vertebrates; see e.g., Whitear, 1989; Brown et al., 2023), Atoh1 acts with Pou4f3 in a conserved ‘feed-forward circuit’, with Atoh1 directly activating *Pou4f3* expression, and Pou4f3 then acting as a pioneer factor to open a significant subset of Atoh1 target enhancers (some shared and some divergent between hair cells and Merkel cells), enabling mechanosensory differentiation (Yu et al., 2021). In hair cells, *Gfi1* is one of the Pou4f3-dependent Atoh1 targets (Yu et al., 2021). Together with the striking conservation of transcription factor gene expression between developing ampullary organs and neuromasts (Modrell et al., 2011b; Modrell et al., 2011a; Modrell et al., 2017b; Modrell et al., 2017a) (also preprint: Minařík et al., 2023), the phenotypes seen in *Atoh1*-targeted F0 sterlet crispant embryos suggest that the molecular mechanisms underlying electroreceptor formation are highly conserved with those underlying hair cell formation. Indeed, the requirement of Atoh1 for *Pou4f3* and *Gfi1* expression in ampullary organs, as well as neuromasts, suggests that the Atoh1-Pou4f3 ‘feedforward circuit’ in mechanosensory cells - i.e., hair cells and epidermal Merkel cells (Yu et al., 2021) - may also be conserved, at least partly, in developing electroreceptors. Taken together, these data support the hypothesis that electroreceptors evolved as a transcriptionally related "sister cell type" to lateral line hair cells (Arendt et al., 2016; Baker and Modrell, 2018; Baker, 2019).

### Electrosensory-restricted Neurod4 is likely redundant with other Neurod family members in sterlet

Paddlefish *Neurod4* was the first-reported ampullary organ-restricted transcription factor gene (Modrell et al., 2017a), with conserved expression in sterlet (this study). We were unable to detect a lateral line organ phenotype in *Neurod4*-targeted sterlet crispants. However, we found that *Neurod1*, *Neurod2* and *Neurod6* are all expressed in sterlet ampullary organs (as well as neuromasts), suggesting that Neurod4 may act redundantly with one or more of these factors in developing ampullary organs. (In paddlefish, however, *Neurod1* expression is restricted to developing lateral line ganglia; Modrell et al., 2011b.) Targeting multiple *Neurod* genes for CRISPR/Cas9-mediated mutagenesis in the future may shed light on the role played by this transcription factor family in ampullary organ development.

### Foxg1 represses electroreceptor formation in the neuromast-forming central zone of lateral line sensory ridges

We also targeted *Foxg1*, a mechanosensory-restricted transcription factor gene that we recently identified in the developing lateral line system of paddlefish and sterlet (preprint: Minařík et al., 2023). *Foxg1* is expressed in the central zones of lateral line sensory ridges where neuromasts form, though excluded from the central domains of neuromasts where hair cells differentiate (preprint: Minařík et al., 2023). Targeting *Foxg1* for CRISPR/Cas9-mediated mutagenesis in F0 sterlet embryos led to a striking phenotype: the formation within neuromast lines of ectopic electroreceptors, often in the large clusters normally seen in ampullary organs. In some cases, ampullary organ fields, which normally flank neuromast lines, effectively ‘merged’ across missing neuromast lines. This phenotype was revealed by examining expression of the electroreceptor-specific marker *Kcnab3*, and two ampullary organ-restricted transcription factor genes: *Mafa* (preprint: Minařík et al., 2023) and *Neurod4* (Modrell et al., 2017a). Thus, *Foxg1* seems to repress an ampullary organ fate within the central domain of lateral line sensory ridges where neuromasts form.

In the mouse inner ear, *Foxg1* is expressed in the prospective cochlea and all sensory patches, in hair cell progenitors and supporting cells (Pauley et al., 2006; Tasdemir-Yilmaz et al., 2021), plus a subset of hair cells (Pauley et al., 2006). Knockout leads to a shorter cochlea with extra rows of hair cells, and to loss or reduction of vestibular end organs (Pauley et al., 2006; Hwang et al., 2009). More hair cells and fewer supporting cells were seen after conditional knockdown of *Foxg1* in neonatal cochlear supporting cells, possibly via the transdifferentiation of supporting cells (Zhang et al., 2019; Zhang et al., 2020). This suggests the possibility that Foxg1 may act in the central zone of lateral line sensory ridges to maintain a proliferative progenitor state, as it does in the mouse olfactory epithelium (Kawauchi et al., 2009).

Furthermore, Fox family members can act as pioneer factors as well as transcription factors (Golson and Kaestner, 2016; Lukoseviciute et al., 2018). A pioneer factor role has been proposed for Foxi3 in otic placode development (see Singh and Groves, 2016). In the developing neural crest, Foxd3 acts early as a pioneer factor, opening enhancers and repositioning nucleosomes to prime genes controlling neural crest specification and migration and, concurrently, to repress the premature differentiation of, e.g., melanocytes (Lukoseviciute et al., 2018). Later in neural crest development, Foxd3 represses enhancers associated with mesenchymal, neuronal and melanocyte lineages (Lukoseviciute et al., 2018). In cortical progenitors, Foxg1 suppresses the adoption at later stages of an early-born cell fate, namely, Cajal-Retzius cells (Hanashima et al., 2004). In the developing chicken otic placode, Foxg1 represses markers of other lineages, such as the olfactory and lens placodes, and epidermis (Anwar et al., 2017). Hence, it is possible that Foxg1 acts in the central zone of lateral line sensory ridges in electroreceptive fishes as a pioneer factor for neuromast/hair cell formation, and/or that it represses ampullary organ/electroreceptor formation. Such a role would be consistent with lateral line expression of *Foxg1* only in electroreceptive fishes: *Foxg1* is not expressed in the developing lateral line of zebrafish or *Xenopus* (e.g., Dirksen and Jamrich, 1995; Papalopulu and Kintner, 1996; Toresson et al., 1998; Eagleson and Dempewolf, 2002; Duggan et al., 2008; Zhao et al., 2009).

Overall, these data lead us to propose the speculative hypothesis that electrosensory organs may be the ‘default’ fate within lateral line sensory ridges in electroreceptive vertebrates, and that Foxg1 represses this fate to enable mechanosensory neuromasts and hair cells to form. To test these hypotheses directly, it will be important in the future to identify global changes in gene expression and chromatin accessibility in the absence of Foxg1.

### Summary and Perspective

Overall, we have found that the ‘hair cell’ transcription factor Atoh1 is required for the formation of lateral line electroreceptors as well as hair cells, consistent with a close developmental relationship between these putative ‘sister cell’ types. Electrosensory-restricted Neurod4 may act redundantly with other Neurod family members expressed in developing ampullary organs. Mechanosensory-restricted Foxg1 represses the formation of electroreceptors within neuromast lines, suggesting the surprising possibility that electroreceptors are the ‘default’ fate within lateral line sensory ridges and raising interesting developmental and evolutionary questions for future investigation.

## Materials and Methods

### Animals

Fertilized sterlet (*Acipenser ruthenus*) eggs were obtained from the breeding facility at the Research Institute of Fish Culture and Hydrobiology, Faculty of Fisheries and Protection of Waters, University of South Bohemia in České Budějovice, Vodňany, Czech Republic, and staged according to Dettlaff et al. (1993). For detailed information about sterlet husbandry, *in vitro* fertilization and the rearing of embryos and yolk-sac larvae, see Stundl et al. (2022). Each fertilization used a mix of sperm from three different males, so each batch was a mix of siblings and half-siblings. Upon reaching the desired developmental stages, embryos and yolk-sac larvae were euthanized by overdose of MS-222 (Sigma-Aldrich) and fixed in modified Carnoy’s fixative (6 volumes 100% ethanol: 3 volumes 37% formaldehyde: 1 volume glacial acetic acid) for 3 hours at room temperature, dehydrated stepwise into 100% ethanol and stored at −20 °C.

All experimental procedures were approved by the Animal Research Committee of the Faculty of Fisheries and Protection of Waters in Vodňany, University of South Bohemia in České Budějovice, Czech Republic, and by the Ministry of Agriculture of the Czech Republic (reference number: MSMT-12550/2016-3). Experimental fish were maintained according to the principles of the European Union (EU) Harmonized Animal Welfare Act of the Czech Republic, and Principles of Laboratory Animal Care and National Laws 246/1992 “Animal Welfare” on the protection of animals.

### CRISPR guide RNA design and synthesis

Target gene sequences were identified using the National Center for Biotechnology Information (NCBI) Basic Local Alignment Search Tool BLAST (https://blast.ncbi.nlm.nih.gov/Blast.cgi; McGinnis and Madden, 2004) to search sterlet transcriptomic data (available at DDBJ/EMBL/GenBank under the accessions GKLU00000000 and GKEF01000000; see preprint: Minařík et al., 2023) or draft genomic sequence data (M.H., unpublished) with the relevant paddlefish sequence (Modrell et al., 2017a). Transcriptomic sequence data were searched for *Tyr*, *Atoh1* and *Neurod4*; genomic sequence data were searched for *Foxg1*. Chromosome-level sterlet genomes became available only after the project started: Du et al. (2020) and the 2022 reference genome (https://www.ncbi.nlm.nih.gov/datasets/genome/GCF_902713425.1/). Open reading frames were identified using the NCBI ORF Finder tool (https://www.ncbi.nlm.nih.gov/orffinder/) and exons annotated by comparison with reference anamniote species (*Lepisosteus oculatus*, *Danio rerio*, *Xenopus tropicalis*) available via Ensembl (https://www.ensembl.org; Cunningham et al., 2022). Conserved domains were identified using NCBI BLASTX (https://blast.ncbi.nlm.nih.gov/Blast.cgi; McGinnis and Madden, 2004). gRNAs were preferentially designed to target 5’ exons, ideally upstream of or within regions encoding known functional domains, to increase the probability of disrupting gene function. gRNAs were designed using the CRISPR Guide RNA Design Tool from Benchling (https://benchling.com) and selected for synthesis based on the following criteria: (1) a high on-target score, ideally no less than 0.5; (2) no off-target matches identified within coding sequences in transcriptome and genome data, unless there were at least two mismatches in the 3’ seed sequence (8-10 bp upstream of the protospacer adjacent motif [PAM], or in the PAM itself); (3) if multiplexing, the gRNA pair were ideally within 50-150 bases of each other to increase the probability of fragment deletion.

DNA templates for CRISPR gRNAs were synthesized using plasmid pX335-U6- Chimeric_BB-CBh-hSpCas9n(D10A) (Addgene, plasmid #42335; Cong et al., 2013) containing the gRNA scaffold. The gRNA scaffold was amplified using a specific forward primer for each gRNA, with an overhang containing the gRNA target sequence and T7 promoter, and a reverse primer that was identical for all reactions. For gRNAs that did not start with G, an additional G was added at the start to ensure efficient transcription. The DNA template was amplified using Q5 polymerase (New England Biolabs, NEB) and purified using the Monarch PCR & DNA Cleanup Kit (NEB). The gRNAs were synthesized using the HiScribe T7 High Yield RNA Synthesis Kit (NEB) and purified using the Monarch RNA Cleanup Kit (NEB) and stored at −80 °C before use. Alternatively, chemically modified synthetic gRNAs were ordered directly from Synthego (CRISPRevolution sgRNA EZ Kit).

### Embryo injection

On the day of injection, 1200 ng gRNA were mixed with 2400 ng Cas9 protein with NLS (PNA Bio) in 5 μl nuclease-free water and incubated for 10 minutes at room temperature to form ribonucleoprotein (RNP) complexes. For gRNA multiplexing, two RNP mixes were combined 1:1 to a final volume of 5 μl, and 0.5 μl of 10% 10,000 MW rhodamine dextran (Invitrogen) added to better visualize the injection mixture and allow selection of properly injected embryos using rhodamine fluorescence. Injection mixtures were kept on ice throughout the injection session. 10 μl glass microcapillaries (Drummond Microcaps) were pulled in a capillary needle puller (PC-10, Narishige) set to 58 °C with two light and one heavy weights, in single-stage pulling mode. Fertilized sterlet eggs were manually dechorionated using Dumont #5 forceps. A 1000 μl pipette tip cut to the same diameter as a dechorionated sterlet egg was used to prepare a series of wells in an agar plate to allow ideal egg positioning during injection using an automatic microinjector (FemtoJet 4x, Eppendorf), set to 100 hPa. Approximately 20 nl of the injection mixture (corresponding to approximately 4.8 ng gRNAs and 9.6 ng Cas9) were injected into fertilized eggs or two-cell stage embryos, targeting the animal pole at a 45° angle. Injected embryos were moved to a clean Petri dish and, for optimum Cas9 efficiency, kept at room temperature until the end of the injecting session or until at least the 32-cell stage, then moved to a 16°C incubator. No more than 30 eggs were kept per 90 mm Petri dish. Unfertilized eggs and dead embryos were removed at the end of the injection day. Petri dishes were checked regularly for dead embryos and the water was changed at least twice a day before gastrulation was completed, and once daily post-gastrulation. Hatched larvae were kept for approximately 16 days post fertilisation until stage 45, then euthanized by MS-222 overdose and fixed with modified Carnoy’s fixative (see above). Fixed larvae were then dehydrated stepwise into 100% ethanol and stored at −20°C.

### Gene cloning, *in situ* hybridization and immunohistochemistry

Total RNA was isolated from sterlet embryos using Trizol (Invitrogen, Thermo Fisher Scientific), following the manufacturer’s protocol, and cDNA synthesized using the Superscript III First Strand Synthesis kit (Invitrogen, Thermo Fisher Scientific). We used our sterlet transcriptome assemblies (from pooled yolk-sac larvae at stages 40-45; preprint: Minařík et al., 2023), which are available at DDBJ/EMBL/GenBank under the accessions GKLU00000000 and GKEF01000000, to design primers for *Neurod4* (forward: GAGAGAGCCCCAAAGAGACGAG; reverse: CTGCTTGAGCGAGAAGTTGACG). cDNA fragments amplified under standard PCR conditions were cloned into the pDrive cloning vector (Qiagen). Individual clones were verified by sequencing (Department of Biochemistry Sequencing Facility, University of Cambridge, UK) and sequence identity verified using NCBI BLAST (https://blast.ncbi.nlm.nih.gov/Blast.cgi; McGinnis and Madden, 2004). For *Neurod1*, *Neurod2*, *Neurod4* and *Neurod6*, synthetic gene fragments based on sterlet transcriptome data, with added M13 forward and reverse primer adaptors, were ordered from Twist Bioscience. GenBank accession numbers are as follows: *Neurod1* OQ808944, *Neurod2* OQ808945, *Neurod4* OQ808946, *Neurod6* OQ808947. The other genes used in this study have been published (preprint: Minařík et al., 2023).

The sterlet riboprobe template sequences were designed before chromosome-level genome assemblies for sterlet were available (Du et al., 2020 and the 2022 reference genome: https://www.ncbi.nlm.nih.gov/datasets/genome/GCF_902713425.1/). Genome analysis showed that an independent whole-genome duplication occurred in the sterlet lineage, from which approximately 70% of ohnologs (i.e., gene paralogs arising from the whole-genome duplication) have been retained (Du et al., 2020). Supplementary File 1 shows the percentage match for each *Neurod* family riboprobe with the two ohnologs, obtained by performing a nucleotide BLAST search against the reference genome (https://www.ncbi.nlm.nih.gov/datasets/genome/GCF_902713425.1/). Equivalent data for the other riboprobes used in this study are available in the preprint Minařík et al. (2023). The percentage match with the ‘targeted’ *Neurod* family ohnolog ranged from 98.9-100%; the percentage match with the second ohnolog was also high, ranging from 96.2-100% (Supplementary File 1), suggesting that transcripts from the second ohnolog are likely to be targeted by each of these riboprobes.

Digoxigenin-labelled riboprobes were synthesized as previously described (preprint: Minařík et al., 2023). Whole-mount *in situ* hybridization (ISH) was performed as previously described (Modrell et al., 2011a). Whole-mount immunostaining post-ISH for Sox2 (rabbit monoclonal, 1:200; ab92494, Abcam) was performed as previously described (Metscher and Müller, 2011), using a horseradish peroxidase-conjugated goat anti-rabbit antibody (1:300, Jackson ImmunoResearch) and the metallographic peroxidase substrate EnzMet kit (Nanoprobes) as per the manufacturer’s instructions.

### Genotyping

To confirm successful mutation in targeted regions, genotyping was performed on trunk and tail tissue that had been removed before ISH and stored in 100% ethanol at −20°C. The tissue was digested using Rapid Extract Lysis Kit (PCR Biosystems) and the target region amplified using HS Taq Mix Red (PCR Biosystems) according to the manufacturer’s instructions. Primers used for genotyping are listed in Supplementary Table 1. After agarose gel electrophoresis, PCR products were extracted using MinElute Gel Extraction Kit (Qiagen) and submitted for sequencing (Genewiz by Azenta Life Sciences). To analyze CRISPR editing efficiency, Sanger trace files were uploaded to the Synthego Inference of CRISPR Edits (ICE) tool (https://ice.synthego.com; Conant et al., 2022).

### Imaging and image processing

Larvae were placed in a slit in an agar-coated Petri dish with PBS and imaged using a Leica MZFLIII dissecting microscope equipped with either a MicroPublisher 5.0 RTV camera (QImaging) controlled by QCapture Pro 6.0 or 7.0 software (QImaging), or a MicroPublisher 6 color CCD camera (Teledyne Photometrics) controlled by Ocular software (Teledyne Photometrics). For most larvae, a stack of images was acquired by manually focusing through the sample, then Helicon Focus software (Helicon Soft Limited) was used for focus stacking. Adobe Photoshop (Adobe Systems Inc.) was used to process images.

## Data availability statement

The publication and associated supplementary figures include representative example images of embryos from each experiment. Additional raw data underlying this publication consist of further images of these and other embryos from each experiment. Public sharing of these images is not cost-efficient but they are available from the corresponding author upon reasonable request.

## Ethics statement

Sterlet animal work was reviewed and approved by The Animal Research Committee of Research Institute of Fish Culture and Hydrobiology, Faculty of Fisheries and Protection of Waters, University of South Bohemia in České Budějovice, Vodňany, Czech Republic and Ministry of Agriculture of the Czech Republic (MSMT-12550/2016-3).

## Author contributions

CB conceived and designed the project and wrote the manuscript together with MM. MM performed most of the experiments, prepared all the manuscript figures and made a significant contribution to the writing of the manuscript. AC and RF performed some experiments. MH provided draft genome sequence data. MP and DG were instrumental in enabling all work with sterlet embryos. All authors read and commented on the manuscript.

## Rights Retention Statement

This work was funded by a grant from the Biotechnology and Biological Sciences Research Council (BBSRC: BB/P001947/1). For the purpose of open access, the author has applied a Creative Commons Attribution (CC BY) licence to any Author Accepted Manuscript version arising.

## Acknowledgements

Thanks to Melinda Modrell for pilot work on CRISPR/Cas9 in paddlefish and for her advice and helpful discussions in planning the project. Thanks to Marek Rodina and Martin Kahanec for their help with sterlet spawns. Thanks to David Jandzik, András Simon, Alberto Joven Araus, Ahmed Elewa, and Mustafa Khokha for invaluable advice on CRISPR/Cas9 approaches and genotyping. Thanks to Jan Stundl and David Jandzik for sharing their sterlet *Tyr* gRNA sequences prior to publication. Thanks to Christine Hirschberger and Rolf Ericsson for their help with some of the *in situ* hybridization rounds.

