## Supplementary Figure_S1-S6 for "Atoh1 is required for the formation of lateral line electroreceptors and hair cells, whereas Foxg1 represses an electrosensory fate"

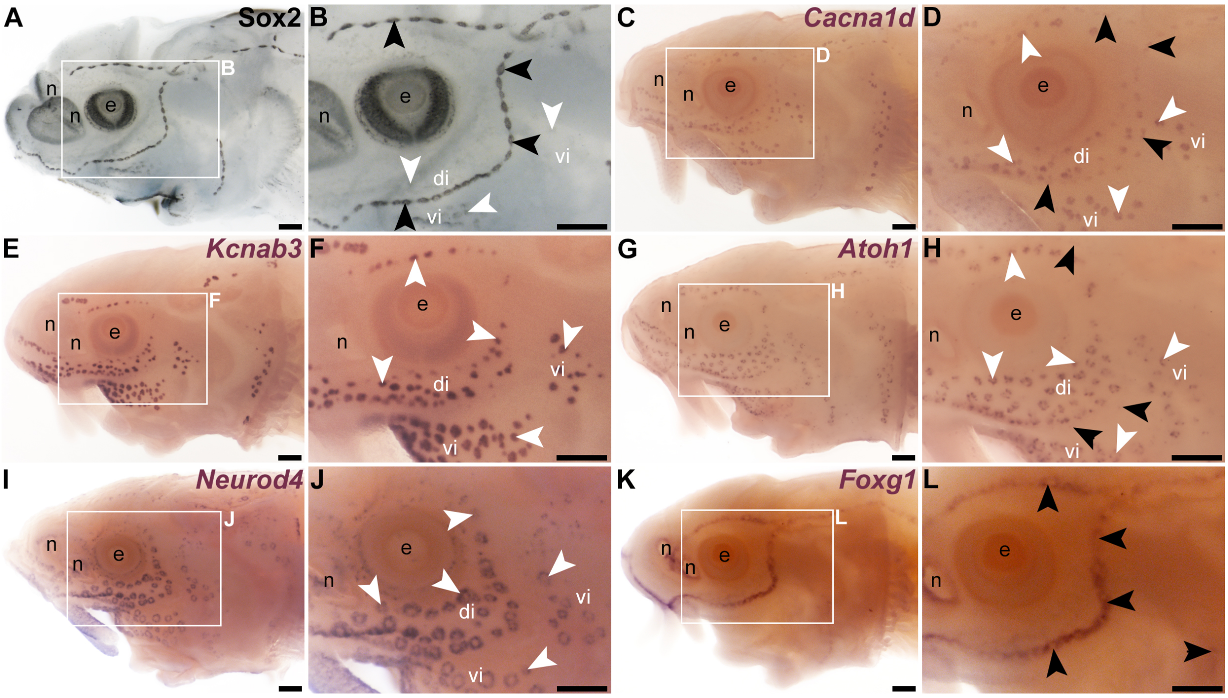


**Supplementary Figure S1. Wild-type gene expression patterns within late-larval sterlet lateral line organs.** Detailed descriptions of the sterlet gene expression patterns shown for reference in this figure, including at earlier stages of lateral line development, have been published (preprint: Minařík et al., 2023), except for *Neurod4*. Sterlet are shown at late yolk-sac larval stages. Black arrowheads indicate examples of neuromasts; white arrowheads indicate examples of ampullary organs. (**A,B**) Immunostaining for the supporting cell marker Sox2 at stage 45 shows strong expression in neuromasts and much weaker expression in ampullary organs. (**C,D**) *In situ* hybridization (ISH) for the hair cell and electroreceptor marker *Cacna1d* at stage 45 shows expression in neuromasts and ampullary organs. (**E,F**) ISH for the electroreceptor marker *Kcnab3* at stage 45 shows expression in ampullary organs only. (**G,H**) ISH for *Atoh1* at stage 42 shows expression in ampullary organs and, more weakly, in neuromasts. (**I,J**) ISH for *Neurod4* at stage 45 shows expression in ampullary organs only. (**K,L**) ISH for *Foxg1* at stage 45 shows expression is restricted to neuromast lines, though excluded from the centres of neuromasts where hair cells form (compare with Sox2 expression in supporting cells in B, and with *Cacna1d* expression in hair cells in D). Abbreviations: di, dorsal infraorbital ampullary organ field; e, eye; n, naris; S, stage; vi, ventral infraorbital ampullary organ field. Scale bar: 200 µm.

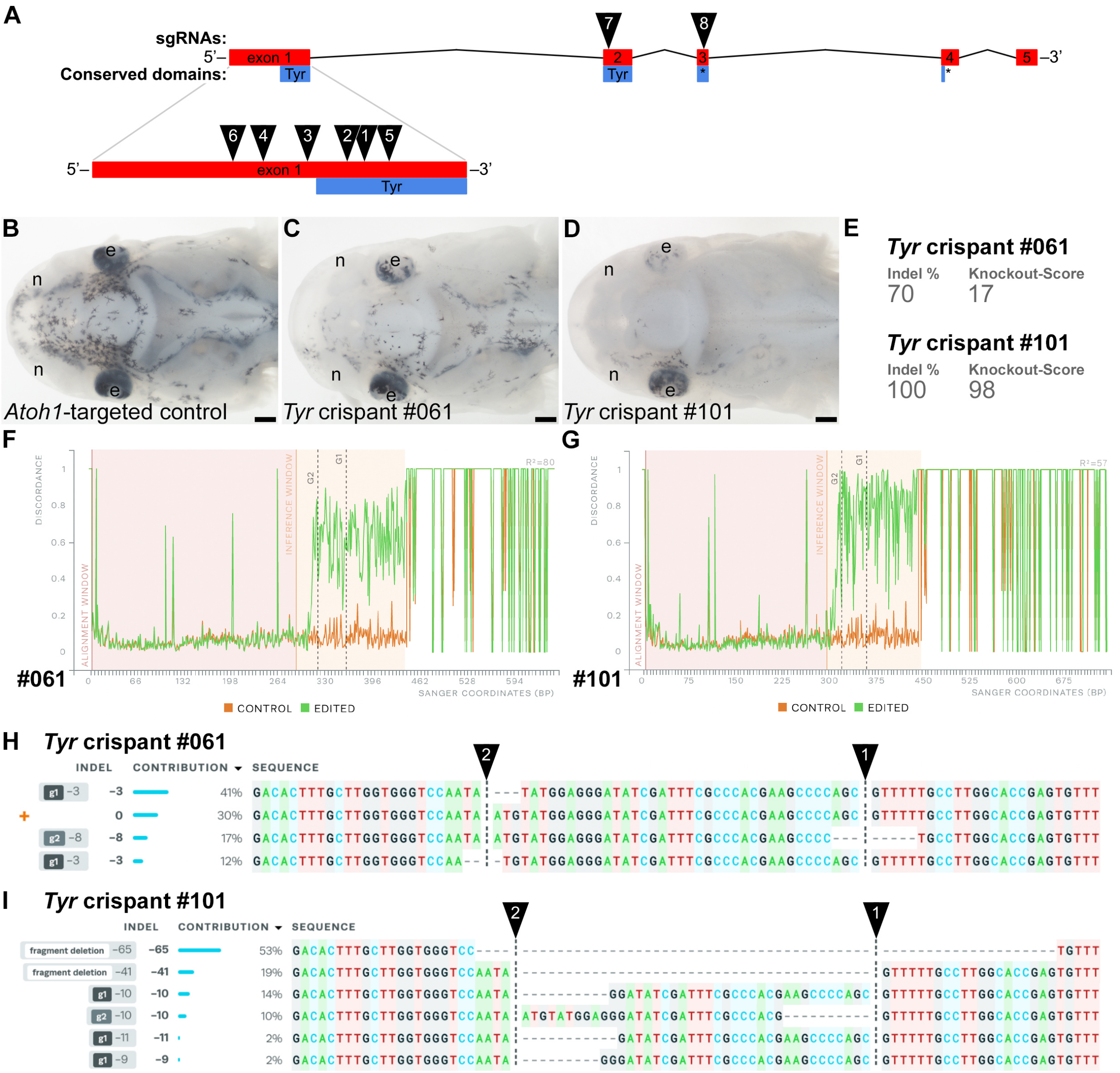


**Supplementary Figure S2. Examples of successful disruption of sterlet *tyrosinase* by CRISPR/Cas-mediated mutagenesis in F0-injected embryos.** (**A**) Schematic showing the exon structure (coding exons only) of the sterlet *tyrosinase* (*Tyr*) gene relative to conserved domains and the target sites of *Tyr* sgRNAs (Table 1). (**B**) Dorsal view of an *Atoh1* crispant at stage 45 as a control (*Atoh1* is not involved in melanin synthesis). Pigmented melanocytes are visible, particularly around the brain, and the eyes are fully pigmented. (**C,D**) Examples of *Tyr* crispants at stage 45 (both targeted with *Tyr* sgRNAs 1 and 2; see A and Table 1) with different degrees of pigment loss (compare with B). In both *Tyr* crispants, significantly fewer melanocytes are visible and the eyes show mosaic loss of pigment. The phenotype is stronger on the right side in both crispants, and stronger in crispant #101 (D) than in crispant #61 (C). (**E-I**) Outputs from Synthego's 'Inference of CRISPR Edits' (ICE) tool (Conant et al., 2022) applied to Sanger sequence data for the targeted region of genomic DNA extracted from the trunk/tail of each of the *Tyr* crispants shown in C and D. 'Indel %' (E) shows the percentage of insertions and/or deletions among the inferred sequences in the CRISPR-edited population. 'Knockout-Score' (E) indicates the proportion of indels that introduce a frameshift or are at least 21 bp long. Discordance plots (F,G) show the level of discordance between the edited sample Sanger trace file (green) and the control sample trace file (orange). Vertical dotted lines indicate the expected cut sites for the sgRNAs. The increase in discordance near the expected cut site indicates a successful CRISPR edit. Nucleotide sequences from the Sanger trace files and their inferred relative contributions to the edited mosaic population are shown in H and I. The expected cut sites are represented by vertical dotted lines. The wild-type sequence (0) is marked by an orange "+" symbol in H, but is absent in panel I because 100% of the sequence was edited in this crispant. Abbreviations: e, eye; n, naris. Scale bar: 200 μm.

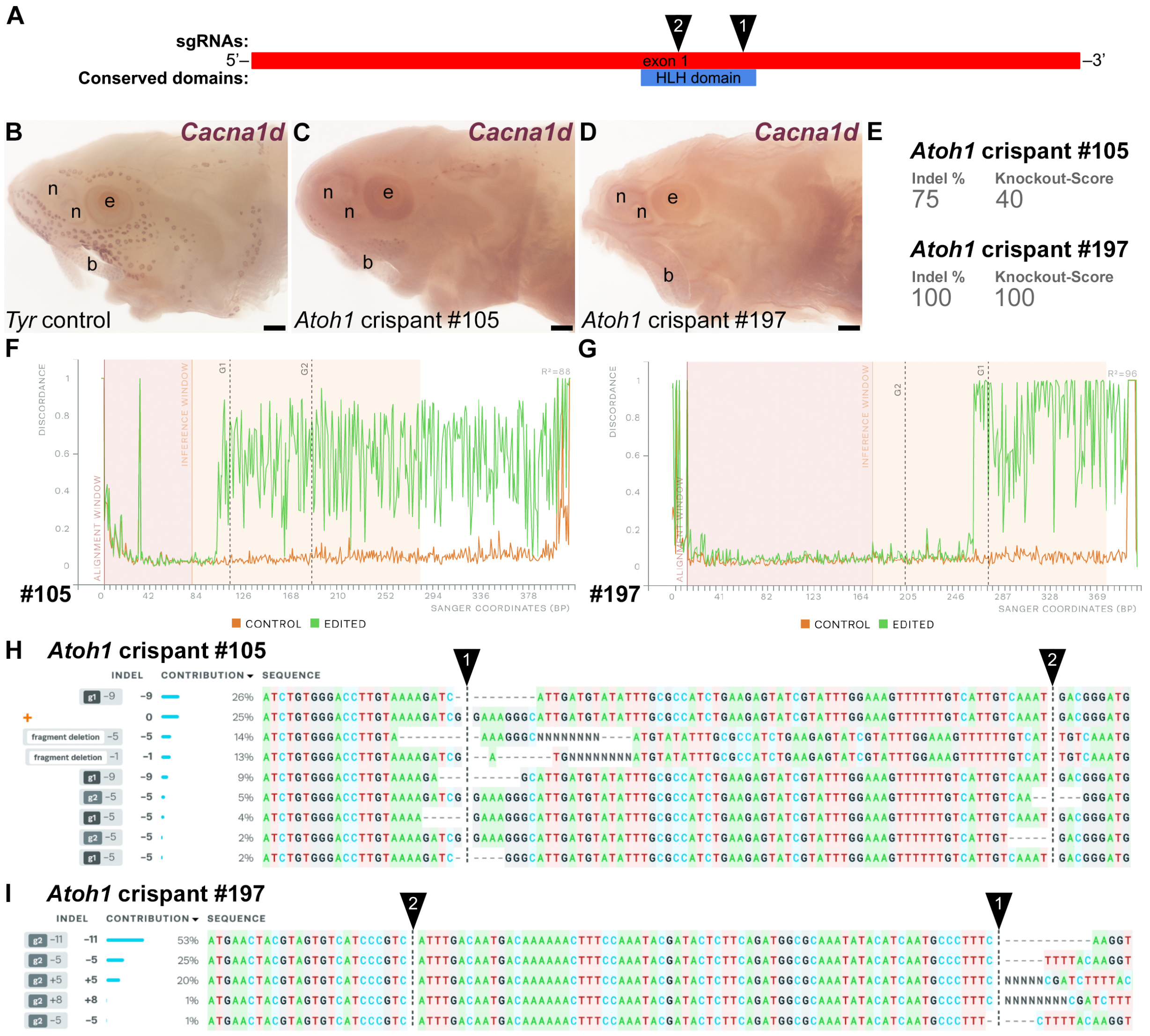


**Supplementary Figure S3. Examples of successful disruption of sterlet *Atoh1* by CRISPR/Cas-mediated mutagenesis in F0-injected embryos.** (**A**) Schematic showing the exon structure (coding exons only) of the sterlet *Atoh1* gene relative to conserved domains and the target sites of *Atoh1* sgRNAs (Table 1). (**B-D**) Sterlet crispants at stage 45 after *in situ* hybridization for the hair cell and electroreceptor marker *Cacna1d*, which is also weakly expressed by taste buds on the barbels. In comparison to the *Tyr* control crispant (B), the two *Atoh1* crispants (C,D: both targeted with *Atoh1* sgRNAs 1 and 2; see A and Table 1) have lost *Cacna1d* expression in neuromasts and ampullary organs, whereas tastebud expression is unaffected. Crispant #105 (C) retains mosaic *Cacna1d* expression in dorsal neuromast lines and a cluster of ampullary organs at the tip of the rostrum. Crispant #197 (D) shows complete loss of *Cacna1d* expression in neuromasts and ampullary organs. (**E-I**) Outputs from Synthego's 'Inference of CRISPR Edits' (ICE) tool (Conant et al., 2022) applied to Sanger sequence data for the targeted region of genomic DNA (chromosome 1 ohnolog) extracted from the trunk/tail of each of the *Atoh1* crispants shown in C and D. 'Indel %' (E) shows the percentage of insertions and/or deletions among the inferred sequences in the CRISPR-edited population. 'Knockout-Score' (E) indicates the proportion of indels that introduce a frameshift or are at least 21 bp long. Discordance plots (F,G) show the level of discordance between the edited sample Sanger trace file (green) and the control sample trace file (orange). Vertical dotted lines indicate the expected cut sites for the sgRNAs. The increase in discordance near the expected cut site indicates a successful CRISPR edit. Nucleotide sequences from the Sanger trace files and their inferred relative contributions to the edited mosaic population are shown in H and I. The expected cut sites are represented by vertical dotted lines. The wild-type sequence (0) is marked by an orange "+" symbol in H, but is absent in panel I because 100% of the sequence was edited in this crispant. Abbreviations: b, barbel; e, eye; n, naris. Scale bar: 200 μm.

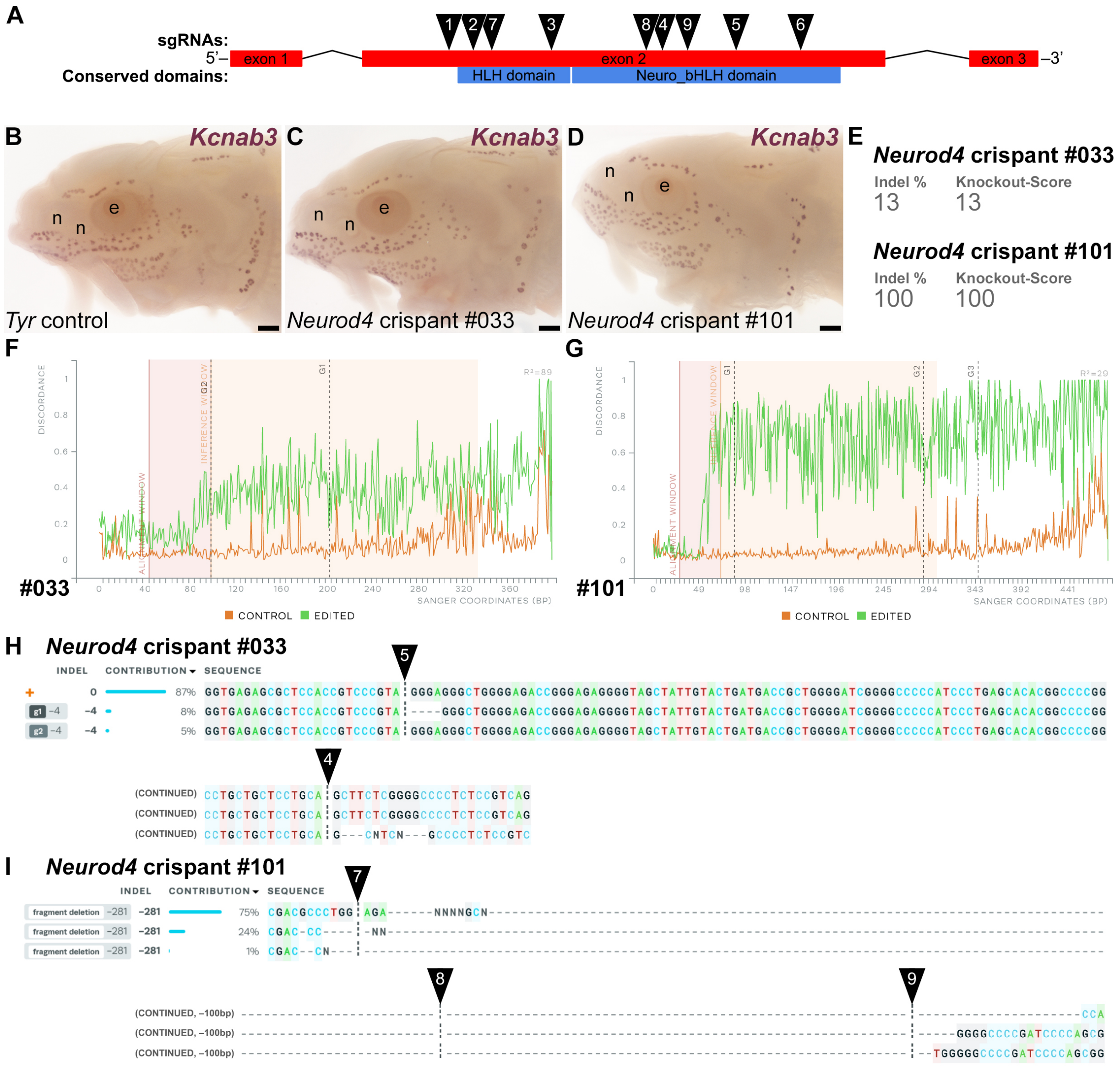


**Supplementary Figure S4. Examples of successful disruption of sterlet *Neurod4* by CRISPR/Cas-mediated mutagenesis in F0-injected embryos.** (**A**) Schematic showing the exon structure (coding exons only) of the sterlet *Neurod4* gene relative to conserved domains and the target sites of *Neurod4* sgRNAs (Table 1). (**B-D**) Sterlet crispants at stage 45 after *in situ* hybridization for the electroreceptor marker *Kcnab3*. In comparison to the *Tyr* control crispant (B), the two *Neurod4* crispants (C,D) show no phenotype. Crispant #033 (C) was targeted with *Neurod4* sgRNAs 3, 4, 5 and 6 (see A and Table 1). Crispant #101 (D) was targeted with *Neurod4* sgRNAs 7, 8 and 9 (see A and Table 1). (**E-I**) Outputs from Synthego's 'Inference of CRISPR Edits' (ICE) tool (Conant et al., 2022) applied to Sanger sequence data for the targeted region of genomic DNA extracted from the trunk/tail of each of the *Neurod4* crispants shown in C and D. For crispant #033, the ICE outputs are shown only for *Neurod4* sgRNAs 4 and 5 (see A and Table 1). 'Indel %' (E) shows the percentage of insertions and/or deletions among the inferred sequences in the CRISPR-edited population. 'Knockout-Score' (E) indicates the proportion of indels that introduce a frameshift or are at least 21 bp long. Discordance plots (F,G) show the level of discordance between the edited sample Sanger trace file (green) and the control sample trace file (orange). Vertical dotted lines indicate the expected cut sites for the sgRNAs. The increase in discordance near the expected cut site indicates a successful CRISPR edit. Nucleotide sequences from the Sanger trace files and their inferred relative contributions to the edited mosaic population are shown in H and I. The expected cut sites are represented by vertical dotted lines. The wild-type sequence (0) is marked by an orange "+" symbol in H, but is absent in panel I because 100% of the sequence was edited in this crispant. Abbreviations: e, eye; n, naris. Scale bar: 200 µm.

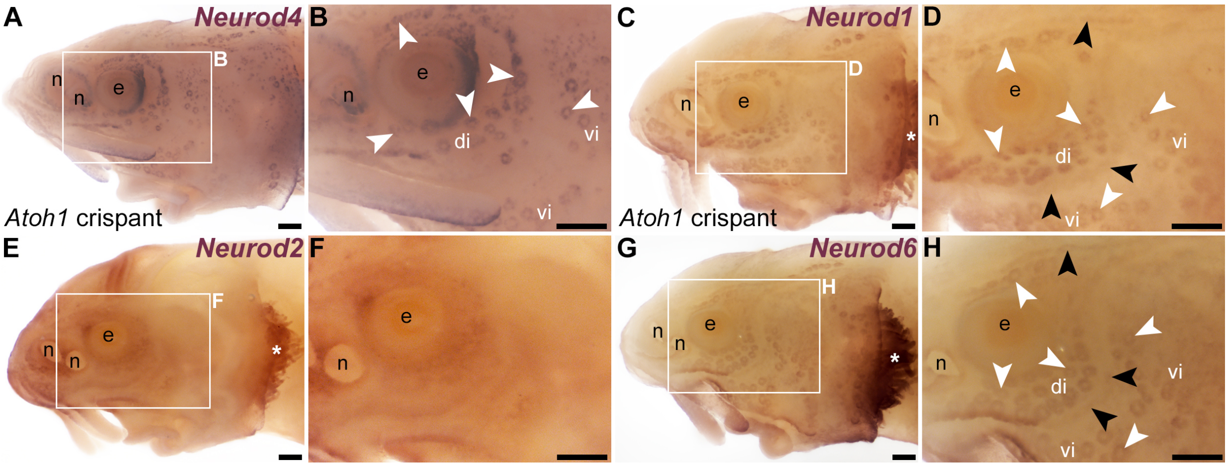


**Supplementary Figure S5. Expression of *Neurod* family genes in late-larval sterlet lateral line organs.** *In situ* hybridization in sterlet at stage 45. Black arrowheads indicate examples of neuromasts; white arrowheads indicate examples of ampullary organs. (**A,B**) *Neurod4* is expressed in ampullary organs only. (**C,D**) *Neurod1* is expressed in ampullary organs and neuromasts (and gill filaments; white asterisk). (**E,F**) No expression was detected in lateral line organs for *Neurod2*, although expression can be seen in gill filaments (white asterisk). (**G,H**) *Neurod6* is expressed in ampullary organs and neuromasts (and gill filaments; white asterisk). Abbreviations: di, dorsal infraorbital ampullary organ field; e, eye; n, naris; vi, ventral infraorbital ampullary organ field. Scale bar: 200 μm.

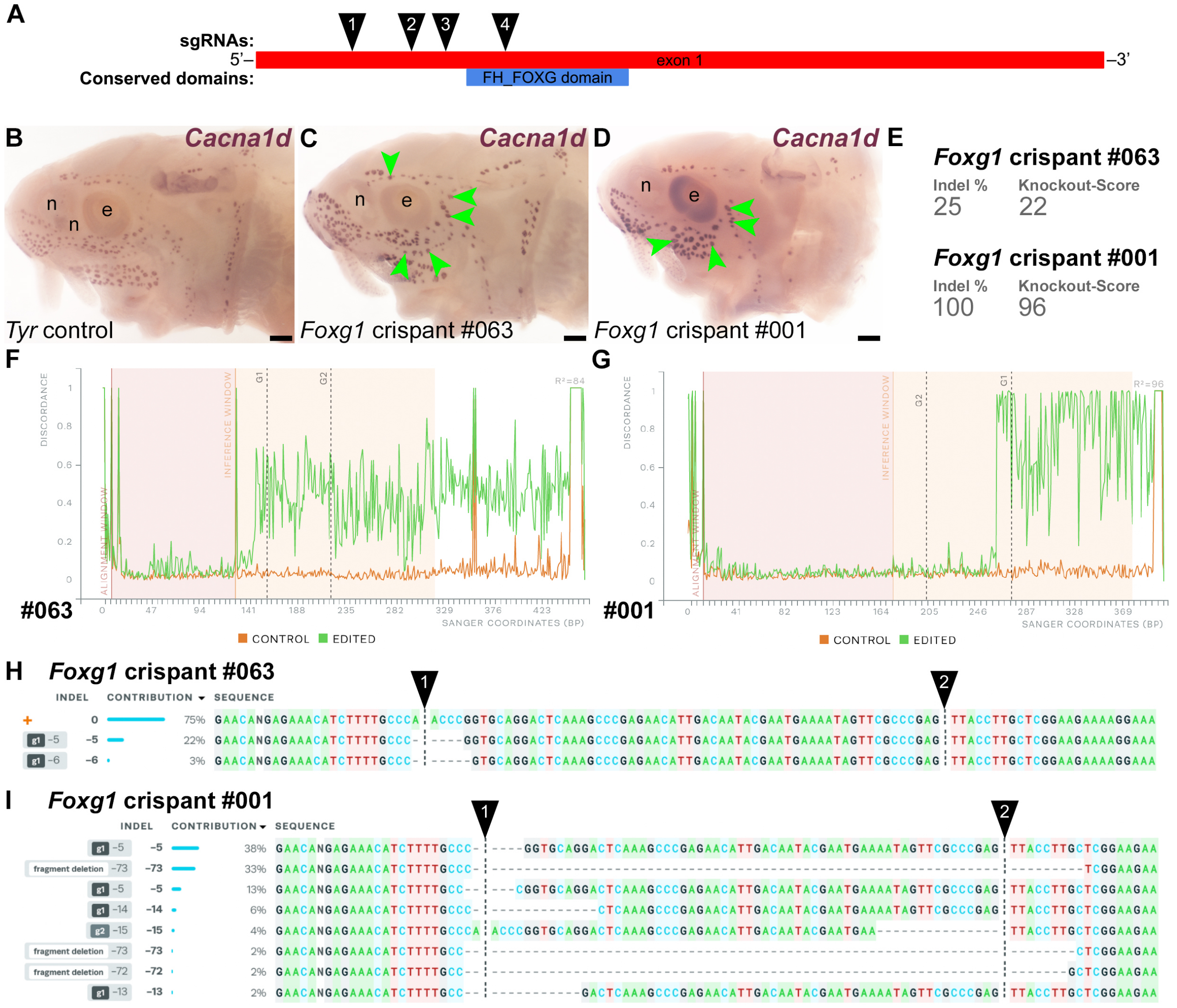


**Supplementary Figure S6. Examples of successful disruption of sterlet *Foxg1* by CRISPR/Cas-mediated mutagenesis in F0-injected embryos.** (**A**) Schematic showing the exon structure (coding exons only) of the sterlet *Foxg1* gene relative to conserved domains and the target sites of *Foxg1* sgRNAs (Table 1). (**B-D**) Sterlet crispants at stage 45 after *in situ* hybridization (ISH) for the hair cell and electroreceptor marker *Cacna1d*. In comparison to the *Tyr* control crispant (B), the two *Foxg1* crispants (C,D: both targeted with *Foxg1* sgRNAs 1 and 2; see A and Table 1) have larger clusters of *Cacna1d*-expressing cells present within neuromast lines than expected for neuromasts, suggesting ectopic ampullary organs (green arrowheads indicate examples). In crispant #001 (D), the infraorbital neuromast line is missing and the ampullary organ fields have fused across it (compare with B and C). (Note: The image in C is also shown in Figure 3I; a ventral view of the crispant in D is shown in Figure 3G.) (**E-I**) Outputs from Synthego's 'Inference of CRISPR Edits' (ICE) tool (Conant et al., 2022) applied to Sanger sequence data for the targeted region of genomic DNA extracted from the trunk/tail of each of the *Foxg1* crispants shown in C and D. 'Indel %' (E) shows the percentage of insertions and/or deletions among the inferred sequences in the CRISPR-edited population. 'Knockout-Score' (E) indicates the proportion of indels that introduce a frameshift or are at least 21 bp long. Discordance plots (F,G) show the level of discordance between the edited sample Sanger trace file (green) and the control sample trace file (orange). Vertical dotted lines indicate the expected cut sites for the sgRNAs. The increase in discordance near the expected cut site indicates a successful CRISPR edit. Nucleotide sequences inferred from the Sanger trace files and their relative contributions to the edited mosaic population are shown in H and I. The expected cut sites are represented by vertical dotted lines. The wild-type sequence (0) is marked by an orange "+" symbol in H, but is absent in panel I because 100% of the sequence was edited in this crispant. Abbreviations: e, eye; n, naris. Scale bar: 200 µm.
